# Local versus global effects of isoflurane anesthesia on visual processing in the fly brain

**DOI:** 10.1101/049460

**Authors:** Dror Cohen, Oressia H. Zalucki, Bruno van Swinderen, Naotsugu Tsuchiya

**Author notes:** Corresponding Authors and, 770, Blackburn Rd, Clayton VIC 3168, St Lucia, Queensland Brain Institute, The University of Queensland, QLD 4072.

## Introduction

Volatile general anesthetics such as isoflurane abolish behavioral responsiveness in all animals, but the neural underpinnings of this phenomenon remain unclear (van Swinderen and Kottler, 2014). Although the cellular and molecular mechanisms through which general anesthetics work have been quite well characterized (Franks, 2008, Garcia et al., 2010, Brown et al., 2011), it is unclear what aspect of neural activity is at the core of the profound disconnection from the environment that is induced by all general anesthetics. To some extent, our understanding of the mechanisms of general anesthesia has paralleled our understanding of how the brain works, from neurons and action potentials, to receptors and neurotransmitters, to circuits and modules, and most recently communication between brain areas. One way to reconcile these varied observations is that general anesthetics might target multiple processes, from sleep circuits to synaptic release (van Swinderen and Kottler, 2014).

General anesthetics have several stereotypic effects on neural activity as measured by the electroencephalogram (EEG). The best known of which is the increase in delta (0.5-4Hz) power that is associated with alternation between highly coordinated UP (depolarized) and DOWN (hyperpolarized) states, which is also observed during non-REM sleep (Lewis et al., 2012, Murphy et al., 2011). EEG studies using propofol anesthesia also show an increase in coherent frontal oscillations in the alpha band (8-12Hz), a potential mechanism for impaired cortical communication (Cimenser et al., 2011, Supp et al., 2011). Studies combining Transcranial Magnetic Stimulation (TMS) with EEG show that midazolam, propofol and xenon dramatically disrupt cortico-cortical communication in response to a TMS pulse (Ferrarelli et al., 2010, Sarasso et al., 2015). These are in agreement with the theoretical suggestion that anesthetics cause the loss of consciousness by interrupting the global integration of cortical activity (Alkire et al., 2008).

The fly model offers a unique opportunity for studying anesthetic action, because it offers the smallest brain (^~^100,000 neurons) that is potentially affected in the same way by general anesthetics as the human brain. Isoflurane anesthesia abolishes behavioral responsiveness in fruit flies (Kottler et al., 2013), and this is associated with decreased brain activity (van Swinderen, 2006). Genetic manipulations in *Drosophila melanogaster* are shedding new light on anesthetic action, suggesting that general anesthesia might also involve presynaptic mechanisms as well as potentiation of sleep circuits (Kottler et al., 2013, van Swinderen and Kottler, 2014, Zalucki et al., 2015). However, it is currently unclear if the effects of general anesthesia on neural processing are conserved across all brains, regardless of specific neuroanatomy. To investigate this we recorded neural activity from multiple regions of the fly brain simultaneously during wakefulness and isoflurane anesthesia, while also measuring brain and behavioral responses to exogenous stimuli.

We used a recently developed multi-electrode preparation (Paulk et al., 2013) to record evoked Local Field Potentials (LFP) across the fly brain in response to flickering visual stimuli. The flickering stimuli produced a periodic response, known as Steady State Visually Evoked Potentials (SSVEPs) (Norcia et al., 2015) that allowed us to accurately track the responses in the frequency domain across brain structures, from the optic lobes to the central brain. We hypothesized that isoflurane would globally reduce the power of the SSVEP throughout the brain, but that impaired signal transmission would have a greater effect in the central brain compared to the optic lobes. We found that isoflurane indeed reduced SSVEP power and coherence in central brain areas but surprisingly responses in the periphery actually increased under isoflurane exposure. We explain these results using a simple model based on known fly neuroanatomy. We further show that the relationship between SSVEP power and coherence can be explained by explicitly considering the relationship between evoked responses and spontaneous brain activity. These results suggest that volatile anesthetics have distinct local and global level effects in all brains, regardless of their specific neuroanatomy, but also that local neuroanatomy is key to understanding anesthetic effects.

## Results

### Evoked responses vary across the fly brain

We presented flickering visual stimuli to awake and anesthetized flies while recording local field potentials (LFPs) from different areas of the fly brain (Figure 1a-c). Before characterizing the effects of anesthesia on the LFPs, we investigated whether different brain areas showed different responses to the visual flicker in 0% isoflurane (air). We first confirmed that in all our recordings (N=16) we were able to detect the SSVEPs, whose magnitude depended on brain region and the location of the visual flicker (Figure 2). When we presented 13Hz flicker ipsilateral (on the same side) to the probe insertion site (Figure 2a), we saw clear periodic waveforms in the time domain (Figure 2b) as well as clear peak at 13Hz and its multiples (harmonics) in the frequency domain (Figure 2c), reflecting robust SSVEPs. Figure 2d summarizes the average SSVEP power at 13Hz (blue)and at its harmonic (26Hz, red), for each channel. SSVEP power at both 13Hz (f_1_) and 26Hz (f_2_) was highest around channel 3-6, roughly corresponding to the medulla of the optic lobe (cyan structure in Figure 1c), and lower responses in channel 8-14, corresponding to the higher-order central structures of the fly brain, as expected and consistent with other SSVEP studies in the fly (Paulk et al., 2015, Paulk et al., 2013).

**Figure 1.**
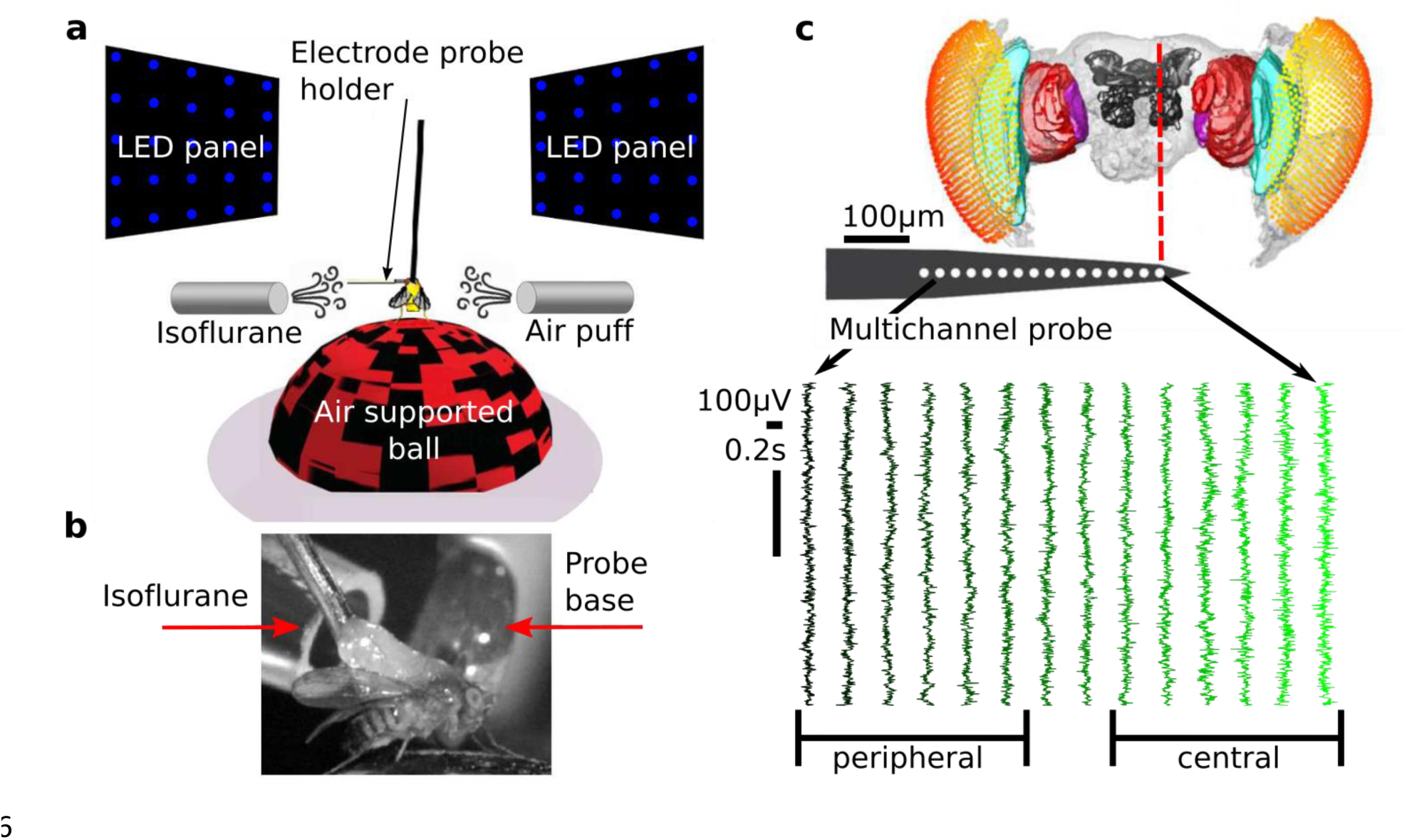
Experimental procedure and paradigm. **a)** Experimental setup. Flies were dorsally fixed to a tungsten rod and placed on an air-supported ball where they could freely walk. Flickering stimuli at 13Hz or 17Hz were presented through two LED screens to the left and right. Isoflurane in different volumetric concentrations was delivered through a rubber hose. An air puff was used as a startle stimulus to gauge the flies’ responsiveness. A 16-contact electrode-probe mounted on an electrode holder was inserted laterally from the left. Only the electrode holder is visible at the depicted scale. **b**) A close up view contralateral to insertion site showing the fly, isoflurane delivery hose and probe base. **c**) Example of spontaneous (no presentation of visual stimuli), bi-polar re-referenced data before anesthesia (0% isoflurane) from a half brain probe recording (see *Electrode probe insertion*). A standardized fly brain is shown for comparison (Paulk et al., 2015, Paulk et al., 2013). The electrode contacts are indicated by white dots (not to scale). Channels are grouped as peripheral, estimated to correspond to the optic lobe, and central, estimated to correspond the central brain.

**Figure 2.**
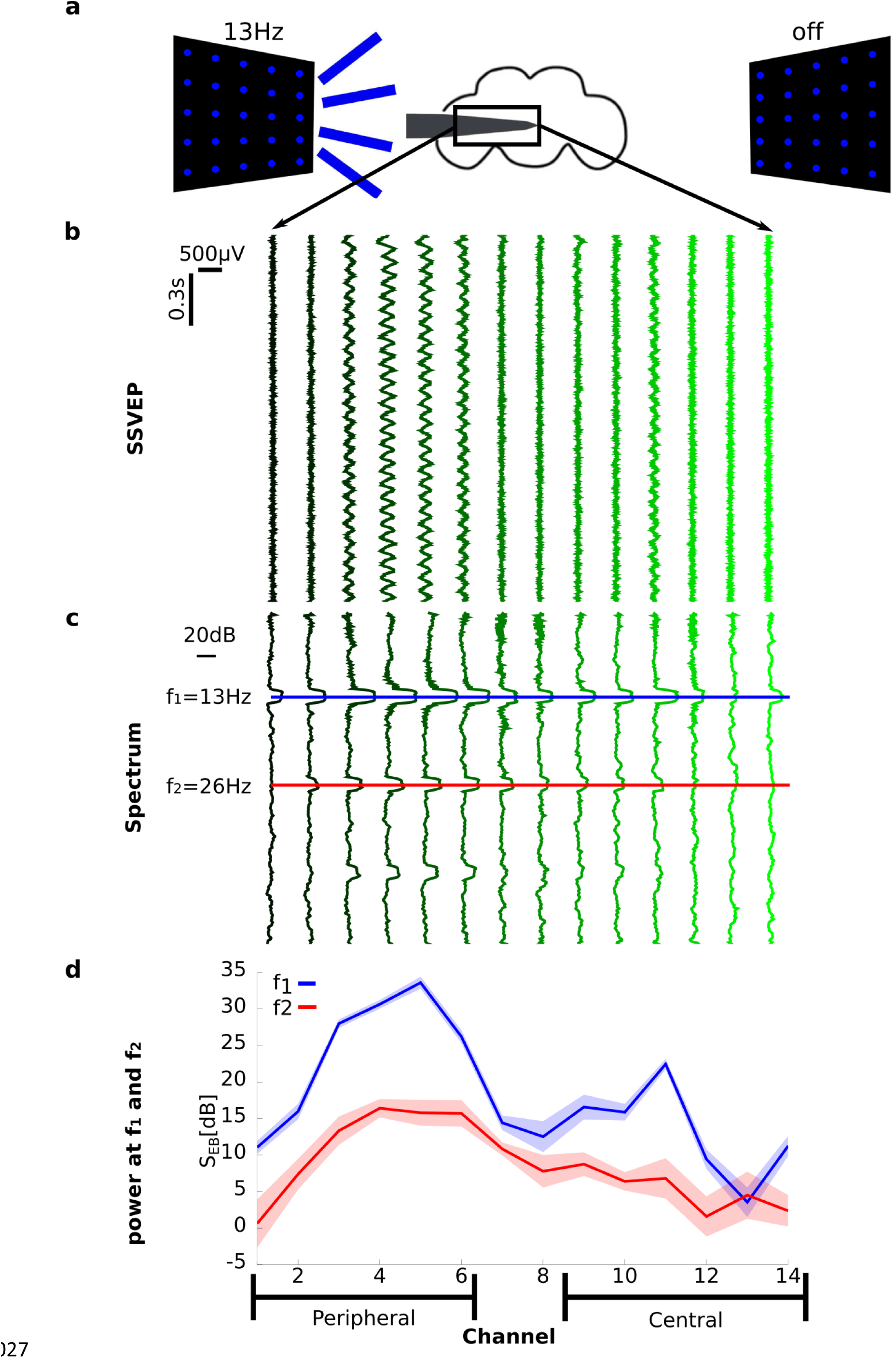
Steady State Visually Evoked Potential (SSVEP) recordings before anesthesia (0% isoflurane). **a**) Schema of an experiment showing the electrode inserted laterally from the left. The left LED panel is shown flickering at 13Hz, corresponding to the [13 off] flicker configuration. **b)** Exemplar mean bipolar re-referenced SSVEP, averaged over 10 trials in the [13 off] condition. The same data from one fly is presented in **c)** and **d)** in different formats. **c)** Exemplar baseline corrected SSVEP power spectrum, averaged over the same 10 trials in **b** (S_EB_(f), see *Local Field Potential analysis*). The blue and the red line mark the first (f_1_=13Hz) and second (f_2_=26Hz) harmonic respectively. **d)** Baseline corrected SSVEP power at f_1_ (blue) and f_2_ (red) for the 10 trials of the [13 off] condition (S_EB_(f_1/2_)). Note narrow shaded areas represent standard deviation across 10 trials, showing the robust and repeatable nature of the SSVEP paradigm. The grouping into peripheral and central channels is depicted at the bottom. The channels are consistently aligned in x-axis **b** through to **d**.

We observed clear and spatially specific responses for both frequency tags (13 and 17Hz; main effect of *channel location*, χ^2^=494.0, p<10^−16^). Classifying flicker configurations as either ipsilateral or contralateral to the insertion site (Figure 3a), we found much higher SSVEP power for ipsilateral flicker configurations than contralateral ones (main effect of *flicker location*: χ^2^=211.0, p<10^−16^), which is in line with the visual information travelling from the optic lobes to the center of the brain (Figure 3b, N=13 flies, 0% isoflurane). For both ipsilateral and contralateral flickers, the responses were stronger at the first (f_1_=13 or 17Hz) than second harmonic (f_2_=26 or 34Hz; main effect of *harmonic*: χ^2^=718.0, p<10^−16^; Figure 3b). In particular, SSVEP power for contralateral flickers at f_2_ was weakest and did not show increased responsiveness in peripheral channels (Figure 3b; confirmed as the significant interaction between *flicker location* and *harmonic*: χ^2^=190.0, p<10^−16^).

**Figure 3.**
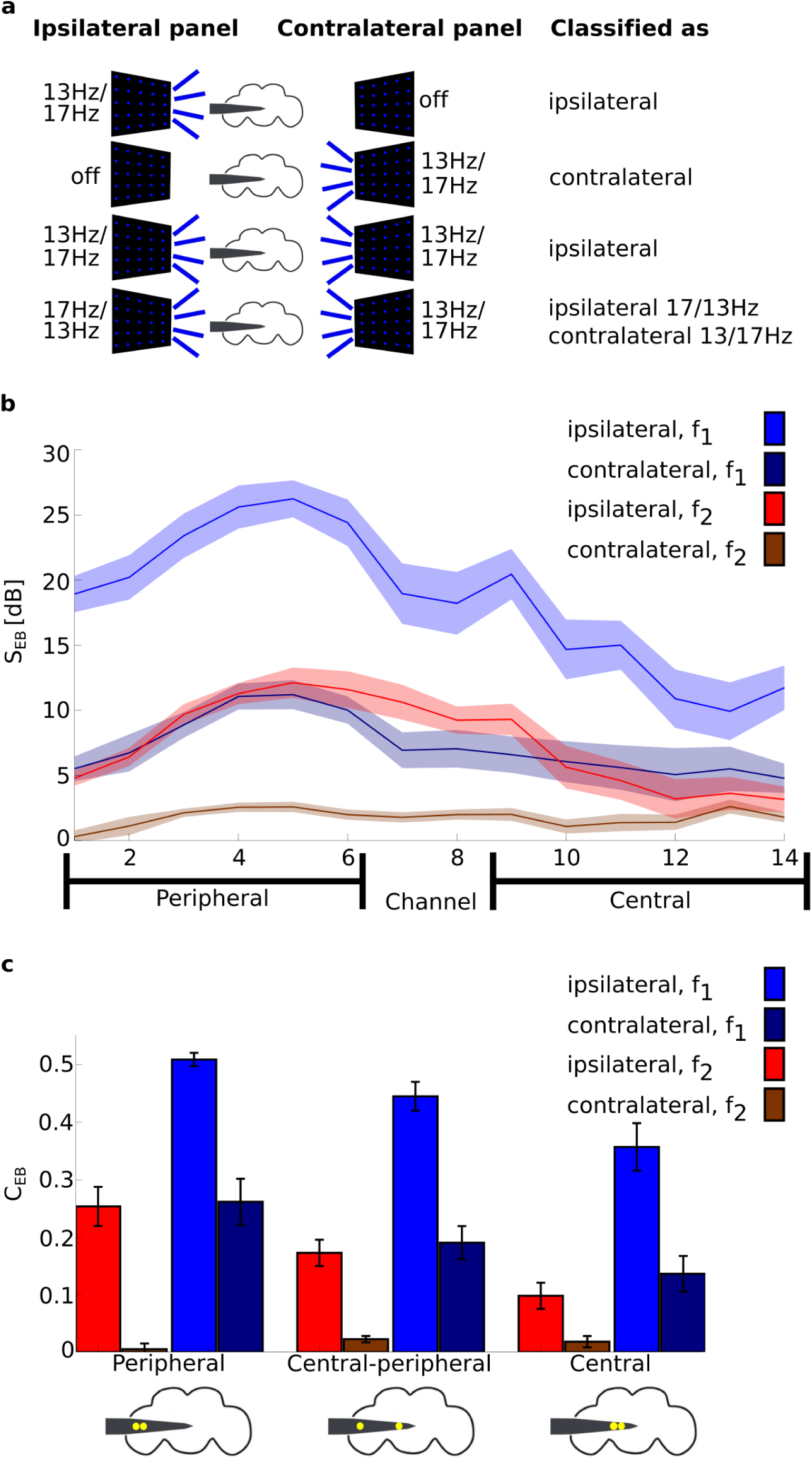
Baseline corrected SSVEP power (S_EB_) and coherence (C_EB_) before anesthesia (0% isoflurane). **a)** Grouping trials according to ipsilateral and contralateral flicker configurations. A trial consisted of a presentation of one of the eight flicker configurations; [off 13], [off 17], [13 off], [17 off], [13 13], [13 17], [17 13] and [17 17]. Trials were classified as either ipsilateral or contralateral according to the location of the flicker with respect to electrode insertion site (rows 1-2). Trials in which the same flicker was presented in both sides were classified according to the flicker that was ipsilateral to insertion (row 3). Trials in which different flickers were presented were classified as ipsilateral at the frequency of the ipsilateral panel and contralateral at the frequency of the contralateral panel (row 4). This way of grouping trials captured much of the variance in the observed neural responses (see *Results*). **b**) Group average (N=13) baseline corrected SSVEP power (S_EB_, see *Local Field Potential analysis*) at f_1_ and f_2_ for ipsilateral and contralateral trials. The SSVEP power was strongest in peripheral channels at f_1_ and attenuated towards the center of the brain. Contralateral flickers still evoked responses, but predominantly at f_1_. Shaded area represents sem across flies. **c**) Group average (N=13) baseline corrected SSVEP coherence (C_EB_) for peripheral (P), central-peripheral (CP) and central (C) channel pairs, as per the grouping in Figure 1b and Figure 2d. Schematics of the fly brain with superimposed examples of channel pairs from each grouping are shown at the bottom. SSVEP coherence followed a similar trend to SSVEP power: higher coherence at f_1_ than f_2_ and a decrease towards the center. Contralateral flickers evoked coherence predominantly at f_1_. Error bars represent sem across flies (N=13).

This observation suggests that the second harmonic, f_2_, reflects more local processing that is evoked mostly when the flicker is presented to the ipsilateral side. In contrast, the first harmonic, f_1_, may reflect more global processing, showing a large response even when the flicker is presented to the opposite side of the insertion site.

### Evoked coherence varies across the fly brain

Next, we assessed whether SSVEP coherence showed brain-region specific patterns. Coherence measures the strength of linear dependency between two variables in the frequency domain (Bendat and Piersol, 2000) and was used in a similar preparation to investigate closed and open loop behavior in flies (Paulk et al., 2015). Paulk et al observed increased SSVEP coherence when flies were engaged in closed-loop behaviour, compared to open-loop, where flies were not in control. Notably, SSVEP power alone did not distinguish between the two conditions.

To summarize the coherence data we grouped channels into periphery and center (Figure 3c) and averaged channel pairs across the periphery, periphery-center and center. Example pairs from each grouping are shown at the bottom of Figure 3c.

Overall, SSVEP coherence showed a similar pattern to SSVEP power. Higher coherence was observed at f_1_ (=13 or 17Hz) than f_2_ (=26 or 34Hz), clearly seen by comparing blue (f_1_) and red (f_2_) bars in Figure 3c (main effect of *harmonic*: χ^2^=179.0, p<10^−16^) and at ipsilateral than contralateral flickers, seen as higher brighter (ipsilateral) than darker (contralateral) bars in Figure 3c (main effect of *flicker location*: χ^2^=29.5, p<10^−6^). The effect of channel location is also strong with the highest coherence observed between peripheral pairs, followed by peripheral-central pairs and weakest for central pairs (Figure 3c; main effect of *channel location*: χ^2^=62.3, p<10^−10^). Similarly to SSVEP power, there was an interaction between *flicker location* and *harmonic* (χ^2^=6.2, p<0.02). While contralateral flicker configurations barely evoked coherent SSVEP activity throughout the brain at f_2_, they evoked location dependent coherence at f_1_. Ipsilateral flicker configurations, however, evoked similar location-dependent coherence at both f_1_ and f_2_ (Figure 3c). Taken together, these similar patterns of results for SSVEP power and SSVEP coherence (Figure 3b and c) suggest their strong relationship.

The results so far have shown that SSVEP power and coherence have a characteristic spatial responses profile with higher values in the peripheral optic lobe than in central regions, and that responses at f_2_ may reflect more local processing as observed in the limited response to contralateral flicker configurations (Figure 3b and c). In what follows we investigated how a volatile general anesthetic, isoflurane, affects these distinct visual responses in the fly brain.

### Isoflurane reduces behavioral responses and attenuates endogenous brain activity

In flies, like other animals, the behavioral effects of general anesthesia are investigated through behavioral responses to noxious stimuli, such as mechanical vibrations (Kottler et al., 2013). In our paradigm, we delivered a series of startling air puffs to the tethered fly exposed to different concentrations of isoflurane (blue rectangles in Figure 4a). To quantify the responses to the air puffs, we analyzed video recordings of the experiments (see *Movement Analysis*). Before any anesthesia (0% isoflurane), flies responded to the air puffs by moving their legs and abdomen, and this was visible as differences in pixel intensities between consecutive frames of the video recording (Figure 4b, left column). 0.6% isoflurane rendered flies completely inert, as evident from small differences between consecutive frames (Figure 4b, right column). After the isoflurane concentration was reset to 0%, flies regained pre-anesthesia responsiveness (Figure 4c): the movement index (MI, see *Movement Analysis*) was significantly below 1 at 0.6% isoflurane (p<0.005, paired two-tailed t-test, dof=12) and not different from 1 at the end of the recovery period (p=0.130). MI was significantly lower during 0.6% isoflurane than after the recovery period (p<0.001). This analysis confirms that isoflurane abolishes behavioural responsiveness in fruit flies, as demonstrated in previous studies (Kottler et al., 2013, van Swinderen, 2006), and also that flies can recover from isoflurane in this preparation.

**Figure 4.**
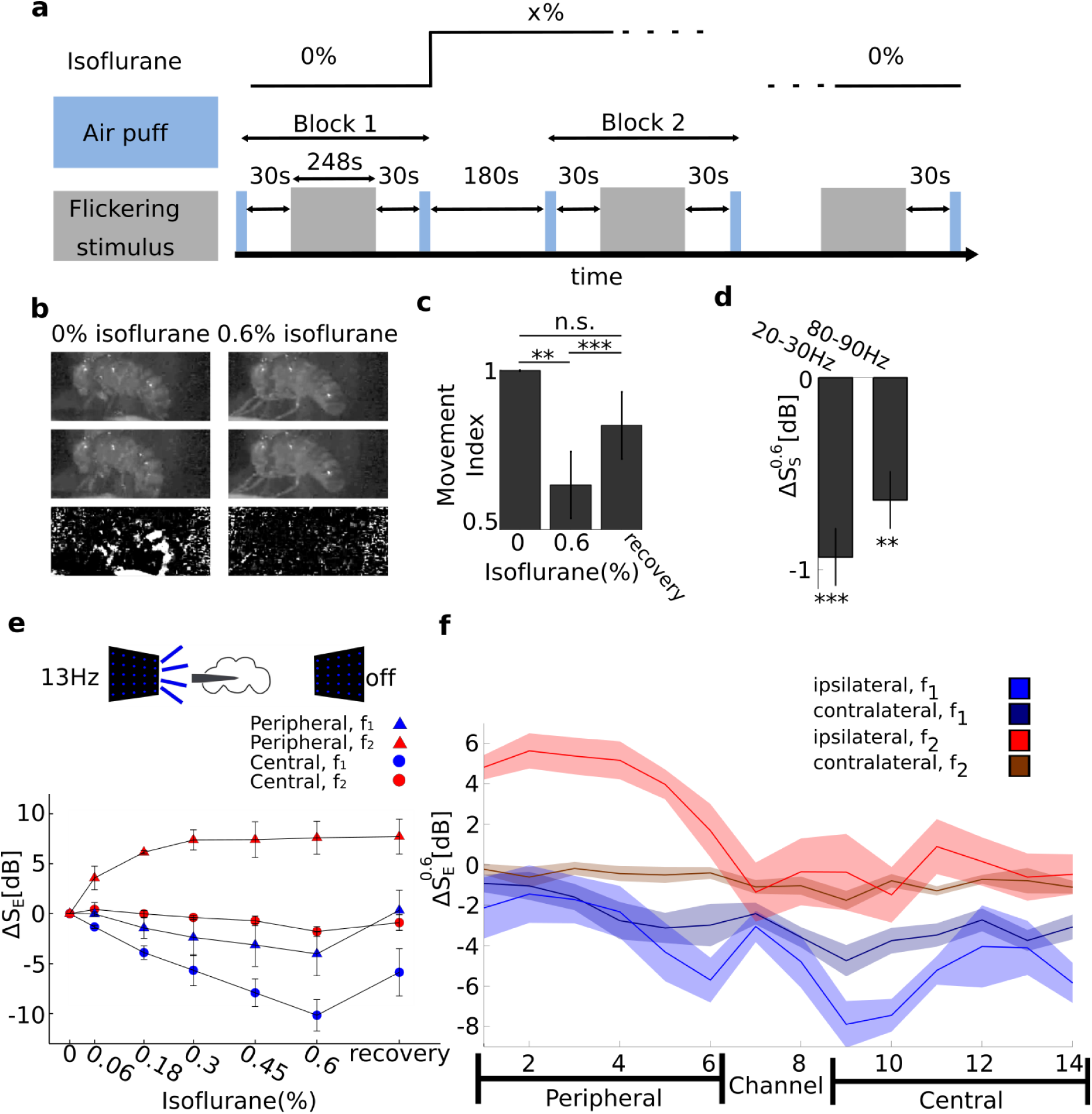
Isoflurane anesthesia has region and harmonic dependent effects on SSVEP power. **a**) Experimental protocol. An experiment consisted of multiple blocks, each at a different concentration of isoflurane (top black line). Each block proceeded with 1) air puffs (light blue rectangles), 2) 30s rest, 3) 80 trials of flicker presentation, corresponding to 10 presentations for each of the eight flicker configurations (gray rectangles), 4) 30s rest, 5) air puff, 6) isoflurane concentration change, and 7) 180s rest for adjustment to the new isoflurane concentration. **b**) Isoflurane abolishes behavioural responses. Consecutive video frames (first and second row) in response to an air puff before any anesthesia (0%, left column) and 0.6% isoflurane exposure (right column). In 0% isoflurane, flies respond to the air puff by moving, seen as the large difference in pixel intensity between consecutive frames (left, third row). In 0.6% isoflurane flies do not respond to the air puff and there are only small differences between consecutive frames. **c**) Quantifying behavioural responses. Group average (N=13) Movement Index (see *Movement Analysis*) was reduced during exposure to 0.6% isoflurane and rebounded after isoflurane levels were reset to 0%. Error bars represent sem across flies. **d)** Isoflurane reduces spontaneous brain activity (ΔS_s_), measured over 4 segments of 2.3s before the start of presentation of the visual stimuli. Group average (N=13) effect of 0.6% isoflurane on spontaneous power (ΔS^0.6^_S_, see *Local Field Potential analysis*). Power is averaged across all channels. Average power for 20-30Hz and 80-90Hz is significantly reduced. Error bars represent sem across flies. **e**) Isoflurane reduces SSVEP power (ΔS_E_) at f_1_ but increases power at f_2_ in a concentration-dependent manner. SSVEP power at f_1_ =13Hz (blue) and f_2_=26Hz (red) for the [13 off] flicker configuration (indicated by the schematic above), at increasing concentrations of isoflurane. For each fly, the SSVEP is first averaged over peripheral (triangles, channel 1-6) or central (circles, channel 9-14) channels. The channel average is further averaged across flies (N=3). Error bars reflect sem across flies. **f**) Isoflurane increases SSVEP power at f_2_ for ipsilateral but not for contralateral flicker configurations. Spatial profile of SSVEP power at f_1_ (blue) and f_2_ (red) for contralateral (dark) and ipsilateral (light) flicker configurations in 0.6% isoflurane (ΔS^0.6^_E_). SSVEP power is averaged across ipsilateral or contralateral flicker configurations (see Figure 3a) first, then averaged across flies (N=13). Shaded area represent sem across flies. The SSVEP power at f1 is reduced in central channels for all flicker configurations, indicating an effect on global neural processing. In contrast SSVEP power at f_2_ is increased at the periphery, but only for ipsilateral flicker configurations, indicating an effect on local neural processing. The peripheral and central channels over which the average was taken in e are depicted at the bottom of f. *** for p<0.001 and ** for p<0.01 in c and d.

Previous work has shown that the attenuated motor behavior in fruit flies is accompanied by attenuated spontaneous brain activity, quantified as a reduction in mean power in the 20-30Hz and 80-90Hz frequency bands (van Swinderen, 2006). We replicated the same effect here using the multiple electrode preparation, by averaging spontaneous power (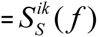, see *Local Field Potential analysis* for the definition) across each frequency band (f) and across all channels (i) at k=0.6% isoflurane concentration. Figure 4d shows the group average (N=13) effect of 0.6% isoflurane on spontaneous power and confirms a significant reduction due to anesthesia in the 20-30Hz and 80-90Hz frequency bands (paired two-tailed t-test, dof=12, p<0.00004 and p<0.001, respectively).

Previous studies in humans show that general anesthetics have spatially distinct effects on spectral power and coherence (Cimenser et al., 2011). In these studies, primary sensory areas tend to remain reliably responsive, but higher-order areas show markedly reduce responsivity (Liu et al., 2012, Mashour, 2013, Supp et al., 2011). Isoflurane is known to potentiate GABAergic neurons, resulting in increased inhibition (Alkire et al., 2008, Garcia et al., 2010) and this is consistent with the attenuated endogenous brain activity observed above and in previous reports (van Swinderen, 2006). Thus, we hypothesized that isoflurane would generally reduce neural responses, but that this reduction would be brain region dependent.

### Isoflurane has opposite effects on SSVEP power at f_1_ and f_2_

To assess the effects of anesthesia on SSVEP power we presented visual flickers during exposure to increasing concentration of isoflurane (gray rectangles in Figure 4a).

In line with our expectations we observed a concentration-dependent reduction in SSVEP power at f_1_. The anesthetic effect was stronger in the central brain than in the periphery (Figure 4e, blue circles). When presenting a 13Hz flickering stimulus ipsilateral to probe insertion site, we found a concentration-dependent reduction that was more pronounced in central than peripheral areas (Figure 4e blue circles versus blue triangles, mean of channels 9-14, N=3). Surprisingly, responses at f_2_ *increased under anesthesia*, but only in peripheral areas (red triangles versus circles in Figure 4e). We confirmed a strong effect for isoflurane concentration (main effect of *isoflurane* (N=3; χ^2^=631.36, p< 10^−16^) as well as an interaction between *harmonic* and *isoflurane* (N=3; χ^2^=434.7, p< 10^−16^)).

To better understand the dissociation between the responses at f_1_ and f_2_, we collected data from 10 additional flies in which we manipulated isoflurane concentration in a binary manner (0% (air) -> 0.6% -> 0% (recovery)). We found that isoflurane reduced SSVEP power at f_1_ (i.e. ΔS_E_ (*ƒ*_1_) < 0) for both ipsilateral (light blue in Figure 4f) and contralateral flicker configurations (dark blue in Figure 4f). In contrast, isoflurane increased SSVEP power at f_2_ (i.e., ΔS_E_(*ƒ*_2_)*>* 0) for peripheral areas but only in response to ipsilateral flicker configurations (light red in Figure 4f). This region-and flicker-specific dissociation was confirmed by a strong interaction between *isoflurane* and *channel location* (χ^2^=187, p<10^−16^) and *isoflurane* and *flicker location* (χ^2^=31.26, p<0.01), as well as the triple interaction between *isoflurane, channel location* and *harmonic* (χ^2^=23.09, p<0.048).

The reduction in SSVEP power at f_1_ was observed for both ipsilateral and contralateral conditions (Figure 4f), suggesting a reduction in global levels of neuronal processing. Further, that this reduction is more pronounced in the central brain is consistent with isoflurane modulating sleep/wake pathways in the central brain (Kottler et al., 2013), in addition to possibly also impairing signal transmission from the periphery to the center. At the same time, the increase in SSVEP power at f_2_ in the periphery was not observed for contralateral flickers, suggesting that this increase may be attributed to isoflurane acting on some local circuit in the periphery. In the following, we provide a potential explanation with simple, yet biologically plausible modeling of the SSVEPs.

### A minimal model explains the opposing effects of isoflurane on SSVEP power at f_1_ and f_2_

A global reduction in SSVEP power at f_1_ is in line with the reduced neural responsiveness described previously (van Swinderen, 2006) and with global impairment of neural communication across the brain (Alkire et al., 2008). However, the local increase in SSVEP power at f_2_ in the periphery is not consistent with these. Here, we propose a minimal model that explains these results in a quantitative manner.

First, we considered what processing of the input can result in a response at f_2_. Linear models of SSVEPs from human electroencephalograms (EEG) have demonstrated reasonable fit to the observed data (see for example (Capilla et al., 2011)). However, we can immediately reject purely linear models because our flickering stimuli consisted of a square wave, whose Fourier decomposition consists of only odd harmonics (f_1_, f_3_, f_5_,…). A linear transformation of the input signal cannot result in power at frequencies which are not present at the input in the first place (Norcia et al., 2015). This suggests that a nonlinear process is involved in the generation of the power at f_2_. What physiological feature in the fly periphery could account for this nonlinearity?

A prominent property in visual processing in animals, including fruit flies, is the segregation of the input pathway into luminance increment responsive (On) and luminance decrement responsive (Off) pathways (Joesch et al., 2010). Splitting of processing into these two pathways is captured by a nonlinearity in the form of half-wave rectification (Regan and Regan, 1988). Half-wave rectification is also implemented in the fly visual system (Reiff et al., 2010) and represents a biologically plausible yet simple nonlinearity.

Figure 5a and b summarize our model which is based on previous models of nonlinear SSVEP generation (e.g. (Regan and Regan, 1988)). First, the input is linearly differentiated to extract points of luminance change before passing through two opposite half-wave rectifiers, corresponding to segregation into On and Off pathways. The result is two pulse trains with the same period as the input stimulus and a time delay of half of the stimulus period. The two pulse trains are separately linearly processed by the On and Off pathways and finally summed to give the recorded response. We estimated the impulse responses of the On and Off pathways for each channel from the response to a 20s 1Hz flicker that was obtained before the main 13/17Hz flicker blocks (Figure 5b)(see *Modeling the SSVEPs*).

**Figure 5.**
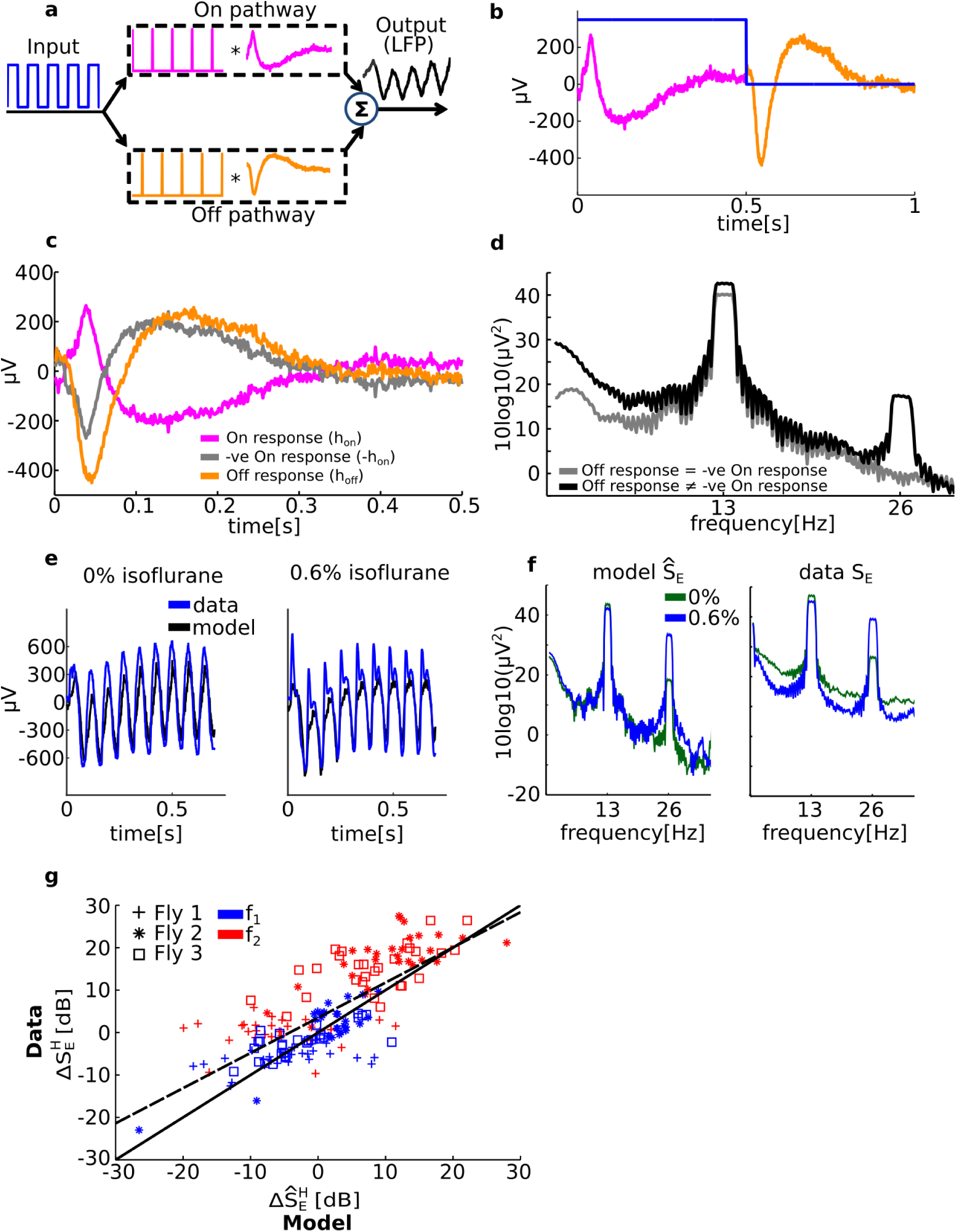
A minimal model explains the unexpected increase in SSVEP power at f_2_ due to isoflurane. **a**) Modeling the SSVEPs. The input (depicted as a blue square wave) is (linearly) differentiated to extract points of luminance increments and decrements before splitting into two streams corresponding to the On (pink) and Off (orange) pathways. Each pathway is modeled as a linear operation determined by the pathway’s impulse response. The responses of the two pathways are summed to give the recorded SSVEP. **b**) The On and Off pathways’ impulse responses were estimated from the response to a 1Hz flickering stimulus (blue). An example from one channel is shown. No other parameters are fitted from the data. **c**) Exemplar On (pink) and Off (orange) impulse responses obtained from the 1Hz flicker presented in both panels [1 1] in 0% isoflurane (air). Note that the negative of the On impulse response (grey) is not identical to the Off impulse response. **d**) The power spectra of the model output. When the negative of the On impulse response is the same as the Off impulse response there is no power at f_2_ (gray). When the empirical On and Off impulse responses are used, the power spectrum has a sharp peak at f_2_ (black). **e)** Comparison between the model output (black) and the recorded SSVEP (blue, average across 10 trials) to a [13 0] stimulus in 0% (left) and 0.6% isoflurane (right) in the time domain. An example from one channel is shown. **f**) Corresponding comparison to **e** in the frequency domain. Spectra of model output (left) and recorded data (right, averaged across 10 trials of the [13 0] flicker configuration) in 0% (green) and 0.6% (blue) show that the model correctly predicts that isoflurane increases SSVEP power at f_2_. **g**) The SSVEP model predictions are in excellent agreement with the observed effects of isoflurane. The model correctly predicts the reduction in power at f_1_ (blue) and increase in power at f_2_ (red), for each of three flies (marked by a cross, square or asterisk), across all channels (14) and both flicker configuration ([13 13] or [17 17]) (n=168, ρ=0.76). The empirical line of best fit (dashed black) closely resembles the line of perfect fit (solid black).

The model predicts that if the Off pathway impulse response is the exact negative of the On pathway impulse response (gray line in Figure 5c), there will be no power at the second harmonic (gray line in Figure 5d). The symmetry between the On and Off responses cancels the nonlinearity (see equation (3.2) in Methods). When the impulse responses for the On and Off pathways are asymmetric, the half-wave rectification is in effect and a prominent peak at f_2_ is observed (Figure 5d, black line).

Our minimal model is effective in explaining the opposing effects of anesthesia at f_1_ and f_2_ in the time (Figure 5e) and frequency (Figure 5f) domain representation of the SSVEP, explaining that isoflurane anesthesia increases the power at f_2_ by changing the impulse responses of the On and Off pathways. We emphasize that the model is completely determined by the response to the 1Hz stimulus, which is then used to predict responses for the [13 13] and [17 17] flicker configurations. No parameters are fitted after computing the impulse responses.

To evaluate the model we computed the correlation coefficient (ρ) and line of best fit between the model-predicted and observed SSVEP power at f_1_ and f_2_ (see *Evaluating the SSVEP model*). We found excellent agreement between the model prediction and the actual data in 0% (air) isoflurane (n=168, ρ^2^=0.9, 95% confidence interval for slope is [1.0 1.21]; for intercept [-6.01 −2.43]) and highest concentration of isoflurane delivered to each fly (n=168, ρ^2^=0.95, slope=[1.05 1.15], intercept=[-1.70 0.75]). Most importantly, the model accurately predicts the *effects* of isoflurane on SSVEP power. Figure 5g shows the observed vs predicted effects of isoflurane on SSVEP power and demonstrates that the model captures both the increase at f_2_ (red) and the decrease at f_1_ (blue) for each of three flies (marked by cross, asterisk and square). The predicted and observed effects of isoflurane show a strong linear relationship (dashed black line, ρ^2^=0.76, dof=167, slope=[0.722 0.94], intercept=[2.50 4.48]) that closely resembles a perfect fit (solid black line).

Thus, according to our model that assumes a minimal yet biologically plausible nonlinearity, the unexpected increase in SSVEP power at f_2_ due to isoflurane anesthesia has a simple explanation; isoflurane changed the balance of the responses of the On and Off pathways. This explanations is also consistent with the observation that the increase was not observed for contralateral flickers, because in this case the On and Off pathways of the *opposite* (i.e., unrecorded) optic lobe would be primarily involved. While our model cannot pinpoint the cellular/molecular mechanisms underlying this change, one potential cause is a wide-spread impairment in synaptic efficacy, independent of sleep circuits, that here results in affected On and Off responses (van Swinderen and Kottler, 2014, Zalucki et al., 2015).

### Isoflurane has opposite effects on SSVEP coherence at f_1_ and f_2_

We next investigated how isoflurane anesthesia affected the observed SSVEP coherence. Generally, the effects were closely related to the changes observed for SSVEP power. Following the same procedure for 0% isoflurane (Figure 3c), we summarized the results by averaging coherence among pairs of recording site within periphery, between periphery and center and within center (see *Analyzing SSVEP coherence*). As expected, 0.6% isoflurane significantly modulated SSVEP coherence (main effect of *isoflurane*, χ^2^=185, p<10^−16^).

The effects of 0.6% isoflurane on SSVEP coherence (ΔC_E_, N=13 flies) are qualitatively similar to those on power, in terms of *channel pair location, flicker location*, and *harmonic* as shown in Figure 6a. Isoflurane reduced coherence at f_1_ but increased coherence at f_2_ (Figure 6a, red vs blue bars, interaction between *harmonic* and *isoflurane*, χ^2^=146, p<10^−16^). The reduction at f_1_ was greater in the center (right column) while the increase at f_2_ was predominantly observed at the periphery (left column, interaction between *isoflurane* and *channel location*, χ^2^=29.25, p<0.00001). The reduction at f_1_ was observed for all flicker configurations (light and dark blue), but the increase at f_2_ was only observed for ipsilateral flicker configurations (light red vs dark red, interaction between *isoflurane* and *flicker location*, χ^2^=12.35, p< 0.009). The triple interaction between *isoflurane, harmonic* and *channel* was not significant (χ^2^=0.66, p=0.72). The results imply that in our paradigm there is a strong connection between SSVEP power and SSVEP coherence, which we dissect in the following section by assuming a linear framework that provides an estimate of coherence based on the Signal to Noise Ratios.

**Figure 6.**
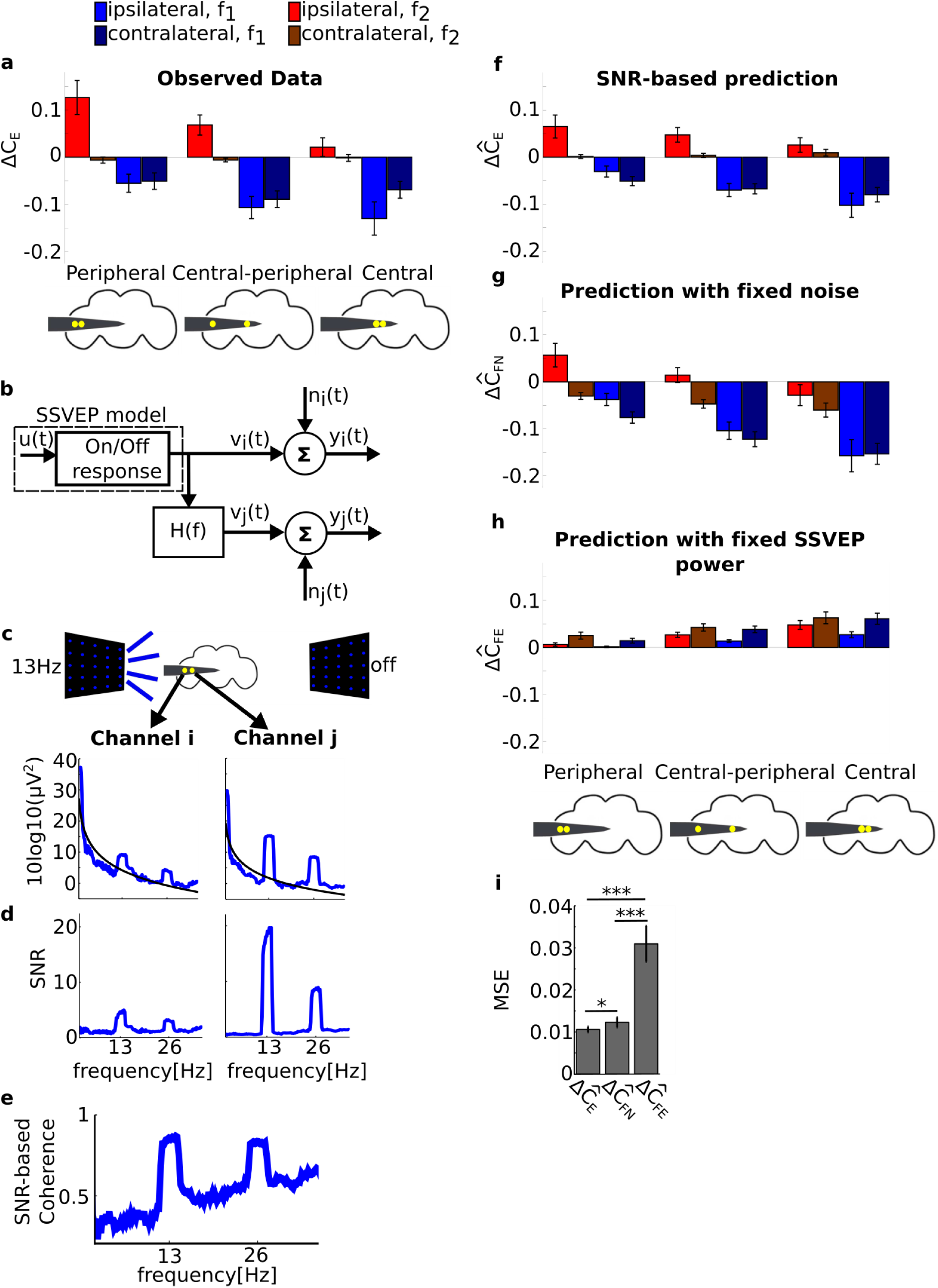
A minimal model based on Signal to Noise Ratios (SNRs) explains the effects of isoflurane on SSVEP coherence. **a**) Group average (N=13) of the observed effects of anesthesia on SSVEP coherence (ΔC_E_). Isoflurane decreases SSVEP coherence at f_1_ (blue) and increases coherence at f_2_ (red). Isoflurane decreases coherence at f_1_ for all flicker configurations throughout the brain. Isoflurane increases SSVEP coherence at f_2_ at the periphery, but only for ipsilateral flicker configurations (light red). Schematics of the fly brain with superimposed examples of channel pairs from each grouping are shown at the bottom. Error bars represents sem across flies. **b**) Linear framework for SNR-based coherence estimates. The SSVEP in channel v_i_(t) is related to the SSVEP in channel v_j_(t) through the transfer function H(f). Independent noise n_i_(t) and n_j_(t) enters at each channel separately to give the recorded SSVEPs y_i_(t) and y_j_(t). Under this scheme, SSVEP coherence has an analytic expression based on the SNR at each channel, given by equation (4.1) (see *SNR-based estimation of coherence*). **c-e)** Estimation of SNR and coherence prediction from the data. **c**) Noise levels were estimated from non-tagged frequencies at each channel, isoflurane concentration and flicker configurations by fitting power-law noise to the SSVEP spectrum (see *SNR-based estimation of coherence*). Exemplar average SSVEP spectra and power-law fits for the [13 off] flicker configuration in 0% isoflurane for two channels, indexed by i and j, are shown. A schematic of the fly brain and channel locations is shown at the top. **d**) The SNR of channels i and j, obtained by dividing the spectrum of the SSVEP by the power-law fit in the linear scale. **e**) Example of SSVEP coherence prediction for channel i and j in panel **d** based on the SNR through equation (4.1). **f**) The SNR-based model correctly predicts the effects of isoflurane on coherence (ΔĈ_*E*_). Group average (N=13) SNR-based prediction of the effects of 0.6% isoflurane. The format and color scheme is the same as in panel **a. g** and **h**) Coherence predictions using different definitions of the SNR. **g**) Prediction based on SNRs using noise levels in 0% isoflurane (ΔĈ_*FN*_, see *SNR-based estimation of coherence*). **h**) Prediction based on SNR using SSVEP power levels in 0% isoflurane (ΔĈ_*FE*_). **i**) Quality of coherence prediction from each model. Mean Square Error (MSE) between the observed (**a**) and each of the three predictions (**f-h**) averaged across all flies, channels, flicker configurations and f_1_ and f_2_. This demonstrates that the effects of isoflurane on SSVEP coherence is largely attributed to the effects of isoflurane on SSVEP power, not on noise. Error bars represent sem across flies (N=13). *** for p<0.001 and * for p<0.05.

### A minimal model explains the opposing effects of isoflurane on SSVEP coherence at f_1_ and f_2_

The observation that isoflurane affected coherence and power in a similar way suggests that in our data the two measures are linked. We explain the opposing effects of isoflurane on SSVEP coherence at f_1_ and f_2_ with another simple model. The model assumes that in each pair of channels, one channel (v_i_(t) in Figure 6b) receives an input from the initial sensory processing (i.e., On/Off response box in Figure 6b) and the other (v_j_(t)) is a linearly filtered version of the first, (represented by the transfer function H(f)). Finally, independent noise (n_i_(t), n_j_(t)) enters at each channel giving the two output voltages (y_i_(t) and y_j_(t)). This simple framework allows us to apply an analytic derivation of coherence based on the Signal to Noise Ratios (SNRs) at each channel (see *SNR-based estimation of coherence* and (Bendat and Piersol, 2000)).

To quantify the SNRs we first estimated the noise level by fitting power-law noise to the power spectrum at the non-tagged frequencies during visual stimulation for each channel, flicker configuration and isoflurane concentration (Figure 6d, see *SNR-based estimation of coherence*). Note that the SSVEP paradigm allows us to operationally regard power at the tagged frequency (f_1_ and f_2_) as “signal” and power at non-tagged frequencies as “noise” (Norcia et al., 2015). Dividing the measured SSVEP power by the estimated noise levels (in the linear scale) provides our estimation of the SNR (Figure 6c-d, see *SNR-based estimation of coherence*). The SNR estimates together with equation (4.1) provide a coherence estimate (Figure 6e). Finally, we separately obtained estimates of the SSVEP coherence in 0% and 0.6% isoflurane to predict the effects of isoflurane on coherence ( ΔĈ_E_) (Figure 6f-h).

The predicted effects of isoflurane on SSVEP coherence is in excellent agreement with the observed data (Figure 6a for the observed coherence and 6f for the model prediction). The model captures the general decrease at f_1_ as well as the increase in periphery coherence at f_2_ for ipsilateral flicker configurations.

In this framework, the effects of isoflurane on noise level as well as SSVEP power both contribute to the SNR-based prediction of coherence. But what is the relative contribution of non-tagged “noise” and tagged “signal” to our successful prediction of SSVEP coherence? To isolate the relative contribution, we re-calculated the SNR by fixing either noise or signal to 0% isoflurane levels, which we call SNR_FN_ and SNR_FE_ (see *Separating the contribution of “noise” and “signal” to the SNR-based estimation of coherence*). The results (Figure 6g and h), clearly show that the contribution of the signal (or evoked response) is much more important for the model prediction.

We formally confirmed the above observation by computing the Mean Squared Error between each SNR-based prediction (ΔĈ_E_, ΔĈ_FN_ and ΔĈ_FE_) and the observed data (ΔĈ_E_) across all channel pairs and flicker configurations (Figure 6i, see *Separating the contribution of “noise” and “signal” to the SNR-based estimation of coherence*). Disregarding the effect of isoflurane on power (ΔĈ_FE_) resulted in considerably worse predictions than ΔĈ_E_ (p<0.0001, dof=12) and ΔĈ_FN_ (p<0.0001, dof = 12). However, disregarding the effects of isoflurane on noise only resulted in slightly (but significantly) worse predictions (ΔĈ_E_ vs ΔĈ_FN_, p<0.040, dof = 12). This means that the observed effects of isoflurane on SSVEP coherence, that is global decrease of coherence at f_1_ and local (peripheral) increase of coherence at f_2_, is largely attributed to the effect of isoflurane on SSVEP power at the stimulus’ tagging frequency, rather than general effects on non-tagged frequencies.

## Discussion

In this paper, we showed that isoflurane has distinct local and global effects on the fruit fly brain. This was made possible by our approach that combines pharmacological manipulation of the states of the brain through anesthetics, perturbation of the neural circuits through periodic visual stimuli, analysis of behaviour and neural data and modeling. Together these components synergistically provide a fuller picture of the effects of isoflurane anesthesia on visual processing, which may generalize to the brains of animals other than flies.

As to the mechanisms of anesthesia, recent studies suggest that reduced cortical communication is at the core of the anesthetic-induced loss of consciousness (Alkire et al., 2008, Mashour, 2013). In particular, increased synchronous activity induced by anesthesia has been suggested to adversely interfere with the communication between brain areas (Sarasso et al., 2014, Supp et al., 2011, Lewis et al., 2012), which may explain the failure of the propagation of evoked responses from primary to higher order areas (Liu et al., 2012, Mashour, 2013, Supp et al., 2011). The volatile general anesthetic isoflurane also abolishes behavioural responses in fruit flies at similar concentrations as required for human anesthesia ((Kottler et al., 2013, van Swinderen, 2006) and here), suggesting that the neural mechanisms through which this anesthetic works may be conserved in most animals. Here we questioned whether isoflurane has distinct effects on local and global processing in the fruit fly brain, and thereby investigate whether an entirely different brain neuroanatomy might reflect similar fundamental effects on neural processing under general anesthesia.

Using a multi-electrode preparation allowed us to record from different brain areas simultaneously, and to assess brain region dependent effects. By presenting flickering visual stimuli we could isolate the neural response in the frequency domain. The frequency decomposition revealed specific effects of anesthesia on the first harmonic (f_1_, 13 or 17Hz) and second harmonic (f_2_, 26 or 34 Hz), which reflected global and local visual processing. Our results show that the reduction in behavioural responses is accompanied by attenuated spontaneous brain activity, and this was also true for the SSVEPs in the central brain, which were reduced for all stimulus configurations, indicating an effect on global neuronal processing at f_1_. In contrast, and to our surprise, local responses at f_2_ in peripheral areas increased, but only for ipsilateral flicker configurations. Modeling the SSVEPs was crucial to understanding this unexpected effect, explaining that the f_2_ power *increase* in the periphery can most likely be attributed to isoflurane-induced changes of the On and Off response pathways in the optic lobes. It is important to note that the brain region dependent effects we have found do not fit with a more simple suppression of neural activity, as this would have resulted in a global and uniform reduction of power. We further showed that the analogous effects of isoflurane on coherence can be explained by explicitly considering how isoflurane affects the tagged brain activity (both f_1_ and f_2_). Overall, the reduction in SSVEP power and coherence in the central brain fits with the view that general anesthetics target inter-area neural communication, impairing the transmission of the visually evoked responses from the optic lobes to central brain structures.

### Evoked and spontaneous activity

The characterization of evoked responses, as opposed to spontaneous activity, through the delivery of a controlled input can reveal additional information about the dynamics of the system. In our experiment, the SSVEPs increased in peripheral areas at f_2_ for specific flicker configurations, revealing a clear difference between the effects of isoflurane on the periphery and center of the fly brain. The use of evoked activity in studying general anesthesia may be particularly important because it allows tracking a stimulus-related neural processes across the brain, potentially making it easier to identify impaired inter-area communication. In SSVEP paradigms, the signal is operationally defined as activity at the tag and its harmonics and this assumption makes it straightforward to define an SNR. This is more difficult with spontaneous activity where “signal” and “noise” cannot be easily separated. Our operational definition of “signal” and “noise” following the tradition of SSVEP studies (Norcia et al., 2015) allowed us to explicitly consider how SSVEP at the tag frequency combine with non-tagged activity (through the quantification of the SNR) to influence coherence. In our data, the effects of isoflurane on SSVEP coherence could be largely attributed to the effects of isoflurane on SSVEP power, as opposed to effects on surrounding, non-stimulus related activity (Figure 6 f-i).

Focusing on neural activity at predefined frequencies, however, is also a limitation of SSVEP paradigms as this only probes the system’s behaviour in a narrow range: the tag and its harmonics. This is particularly important in the context of nonlinear systems whose frequency response can be highly input dependent. We expect that both our modeling of the SSVEPs and the SNR based estimation of coherence will need to be expanded when the system is evaluated over a broader dynamic range.

### Neural substrate of the SSVEPs

Our modeling of the SSVEPs concisely yet plausibly accounts for the unexpected increase in power at f_2_ observed in the periphery. Given the vast literature on elementary motion detection circuitry in flies (Borst and Euler, 2011, Egelhaaf and Borst, 1989, Reisenman et al., 2003), it may be possible to provide more comprehensive modeling. However, for the purpose of explaining the unexpected effects of anesthesia, our minimal modeling was sufficient and provided a physiologically plausible explanation; isoflurane most likely affected the responses of local On and Off pathways which, combined with the presentation of a periodic stimuli, resulted in increased power at f_2_.

Even for our simple model it is not straightforward to assign a fine neural substrate to the SSVEP because there are many connections between the fly optic lobes, such that stimulation in one lobe causes activation in the other (Haag and Borst, 2008). While our recordings (and (Paulk et al., 2015)) clearly show that SSVEP power is much smaller when the flicker is presented to the opposite eye, the broad-field flicker prevents us from precisely disentangling the relative contribution of each optic lobe to the LFP. Another factor is the aggregate nature of the LFP; while the first On and Off responsive cells may be observed as early as the lamina (Reiff et al., 2010), we cannot tell how much these cells contribute to the LFP, compared to other downstream neurons. Future studies separating the contributions of the On or Off pathways to the LFP via genetic manipulations and the use of stimuli that target each pathway separately will help clarify neural substrate of the SSVEP.

### Slow wave and inter-area neural communication

Sleep and general anesthesia are defined by similar criteria and there is evidence for some shared mechanisms (Franks, 2008). The involvement of sleep mechanisms in the impairment of cortical communication observed in general anesthesia (Ferrarelli et al., 2010, Sarasso et al., 2015) is not established, but one possibility is that the stereotypical DOWN states that manifest as the EEG slow wave and observed in both general anesthesia and non-REM sleep may prevent long-range coordinated activation (Sarasso et al., 2014).

Recent findings extend the proposed relationship between sleep and anesthesia to fruit flies where genetic manipulations of sleep circuits can confer both resistance and hypersensitivity to isoflurane (Kottler et al., 2013). However, to date, no evidence or analogue to the slow wave has been observed in sleep or anesthesia in flies (Kirszenblat and van Swinderen, 2015, van Swinderen, 2006), and we found no evidence of it here. Thus, anesthetics may target sleep circuits in all brains but only produce a slow wave in some. Instead, the mechanism for the reduced responsiveness in the central brain that we observed under isoflurane may be a combination of potentiated sleep circuits and compromised synaptic efficacy, which has been demonstrated in flies (Zalucki et al., 2015, van Swinderen and Kottler, 2014). While sleep circuits seem unlikely to modulate the responses of the peripheral On and Off pathways, the globally compromised synaptic efficacy could cause an imbalance in the responses of the On and Off pathways resulting in the unexpected increase in power at f_2_.

### Outlook

Bottom-up approaches that focus on molecular mechanisms have considerably improved our understanding of anesthetic drugs, and have identified a promising set of potential target sites (Brown et al., 2011, Franks, 2008, Garcia et al., 2010). On the other hand, it remains unclear how effects at the molecular level affect large-scale neuronal circuits. Instead, top-down approaches that focus on global effects are providing evidence that general anesthetics share a common endpoint in the reduction of inter-area communication (Lee et al., 2013, Mashour, 2014, Sarasso et al., 2015). Using the metrics developed for characterizing these global effects (Casali et al., 2013, Lee et al., 2015) in conjunction with the genetic manipulations available in *Drosophila* is a promising direction. Studies that manipulate the state of the brain and external perturbations can be combined with signal processing techniques and modeling to help us understand how anesthetic effects at the molecular level change the global state of the brain.

## Methods

### Animals

Female laboratory-reared *Drosophila melanogaster* (Canton S wild type) flies (3-7 days past eclosion) were collected under cold anesthesia and positioned for tethering. Following a procedure previously described for this preparation (Paulk et al., 2013), flies were dorsally glued to a tungsten rod using dental cement (Synergy D6 FLOW A3.5/B3, Coltene Whaledent, Altstätten, Switzerland) which was cured with blue light. Dental cement was applied to the neck to stabilize the head. Flies’ wings were glued to the tungsten rod to prevent wingbeats or attempted flight during recording. Tethered flies were positioned above a 45.5mg air-supported Styrofoam ball (Figure 1a,b), similar to that described in (Paulk et al., 2013).

### Electrode probe insertion

Probe insertion was similar to the procedure outlined in (Paulk et al., 2013). Briefly, linear silicon probes with 16 electrodes (Neuronexus Technologies, Ann Arbor, Michigan) were inserted laterally to the flies’ eye and perpendicularly to the eye’s curvature. Insertion was performed with the aid of a micromanipulator (Merzhauser, Wetzlar, Germany), with the electrode recording sites facing posteriorly. For the majority of experiments (14 flies) probes with electrode-site separation of 25μm (3mm-25-177) and 375μm from base to tip (Figure 1c) were used. This probe covers approximately half of the brain and is referred to as the ‘half’ brain probe henceforth. In two additional flies, probe 3mm-50-177, with electrode-site separation of 50μm and measuring 703μm from base to tip was used. This probe covers approximately the whole brain and is referred to as the ‘whole’ brain probe. Probe tip width (33μm), base width (123μm), thickness (15μm) and electrode site area (177um^2^) are identical for both probes.

A sharpened fine tungsten wire (0.01 inch diameter, A-M Systems, Carlsborg, Washington) acted as the reference electrode and was placed superficially in the thorax (Paulk et al., 2013). Recordings were made using a Tucker-Davis Technologies multichannel data acquisition system with a sampling rate of 25kHz (Tucker-Davis Technologies, US).

The probes were fully inserted until all electrode sites were recording neural activity, confirmed by the presentation of visually flickering stimuli (1Hz and 13Hz, see *Visual Stimuli*) and observing SSVEPs at the most peripheral electrode site (furthest from the probe tip). The probe was then gently retracted until the most peripheral site showed little to no neural activity. We assumed that this indicated that the most peripheral site was placed just outside the eye. This ensured consistent probe insertion depth among flies.

### Visual stimuli

Flickering blue lights (spectral peak at 470nm with 30nm half-peak width) were presented through two LED panels (Figure 1a). The panels were flickered on and off (square wave, 50% duty cycle) at approximately 13.4Hz (hereafter 13Hz) or 16.6Hz (hereafter 17Hz), and either to the left or to the right of the fly. There were thus 8 possible flicker configurations. These are [off 13], [off 17], [13 off], [17 off], [13 13], [13 17], [17 13] and [17 17], where the number represents the flicker frequency and the location represents the left or right LED panel. An ‘off’ signifies that we turned off the blue LED lights in the respective panel. Visual flickers were presented in sets of 80 trials, consisting of 10 presentation of each flicker configuration. A trial lasted 2.3s and the inter-trial interval was 0.8s, taking 248s to complete the 80 trials. The flicker configuration order was randomly generated with the added restriction that consecutive trials consisted of different flicker configurations. Panel voltage levels were recorded at 25 kHz in the same recording system as the electrophysiological signals. The LED lights were turned off except during the period of visual stimulation (Figure 4a).

In three of the flies (2 with the whole-and 1 with the half-brain probe, all with the graded anesthesia manipulation, see *Isoflurane delivery*), we also included an additional flicker configuration of [1 1] (1Hz flicker in both panels). This stimulus was presented once for 20s before the start of the 80 trials described above. These three flies are used for evaluating the modelled SSVEPs (see *Modeling the SSVEPs*)

### Grouping flicker configurations as ipsilateral or contralateral

The 8 flicker configurations were chosen to isolate the effects of flicker frequency (13 vs 17Hz), flicker interaction (e.g. [13 off] vs [13 17]) and flicker location (e.g. [13 off] vs [off 13]). However, we found that grouping trials as either ipsi-or contralateral simplified the results and sufficient for all our claims (see *Results*). Under this classification scheme, trials in which a single flicker was presented at the left panel were labelled ipsilateral ([13 off] and [17 off] in Figure 3a row 1). Trials in which a single flicker was presented at the right panel were labelled contralateral ([off 13] and [off 17] in Figure 3a row 2). Because the ipsilateral flicker dominated the response, we classified trials in which the same flicker was presented in both panels as ipsilateral (Figure 3a row 3) and trials in which different flickers were presented in both panels as ipsilateral at the frequency of the ipsilateral panel and contralateral at the frequency of the contralateral panel (Figure 3a row 4).

### Isoflurane delivery

Isoflurane was delivered onto the fly through a rubber hose connected to an evaporator (Mediquip, Brisbane, Australia) (Figure 1a,b). The isoflurane was blown onto the fly at a constant flow of 2l/min and continuously vacuumed from the opposite side of the fly. Following the gas chromatography procedure described in (Kottler et al., 2013) for measuring isoflurane concentration, we found that the actual concentration near the fly body was 0.3% (vol.) when the concentration at the evaporator was set to 1%. Throughout the paper, we report isoflurane concentration as the linearly estimated concentration at the fly body, not at the evaporator.

Isoflurane concentrations were manipulated in either a graded or a binary manner over the blocks. In the graded manipulation (N=3 with the half-brain probe and N=2 with the whole-brain probe), concentrations were incrementally and sequentially increased over 5 levels and then reduced to 0%; 0% (air)→ 0.06%→ 0.18%→ 0.3%→0.45%→0.6%→0%(recovery). In the binary manipulation (N=10 with the half-brain probe), isoflurane concentration was manipulated over 3 blocks; 0% (air) →0.6%→ 0% (recovery). Throughout the paper, we distinguish two periods of 0% isoflurane as 0% (air) and 0% (recovery), before and after drug exposure respectively. In one fly in which we used the half brain probe we administered the graded manipulation up to 0.45%. In a subset of the flies (N=8 out of 14 with the half-brain probe) an additional recovery block, 0% (recovery 2), was performed.

### Air puff stimuli and behavioural responsiveness

An olfactory stimulus controller (custom built, (Paulk et al., 2013)) was used to deliver six air puffs to gauge the flies’ behavioral responsiveness in each concentration of isoflurane. The inter-air puff duration was approximately 1.5s. Air puffs were delivered before and after the presentation of visual stimuli (Figure 4a). Fly movement activity was recorded with 602f-2 Basler firewire camera (Basler, Ahrensburg, Germany) and a 1-6010 Navitar 12x Zoom lens (Navitar, Rochester, New York) at 30 frames per second, time-locked to the onset of the air puff. We used the video data to assess the flies’ behavioral responsiveness under anesthesia (see *Movement Analysis*).

### Experimental protocol

After inserting a probe and confirming flies’ visible responses to an air puff, we initiated our experimental protocol. An experiment consisted of several blocks, each at a different concentration of isoflurane (Figure 4a). Each block started with the delivery of a series of air puffs, used to gauge the fly’s responses and establish the depth of anesthesia (see *Movement Analysis*). 30s after the startle stimulus, 80 trials of visual flickers were presented (see *Visual Stimuli*). After the completion of 80 trials, the flies were left for an additional 30s, and then a second episode of air puffs was delivered. After the last air puff was delivered the isoflurane concentration was immediately changed. Flies were left for 180s to adjust to the new isoflurane concentration before the next block commenced.

### Movement analysis

To confirm the depth of anesthesia, flies’ movements were analyzed in response to the air puffs. The recorded movies were analyzed to extract the amount of overall movement using custom software written in MATLAB (MathWorks, Natick, Massachusetts). First, movies were down-sampled to 5 frames per second and converted to grayscale. Second, individual images were annotated with the corresponding isoflurane concentration (*k*=[0 (air), 0.06, 0.18, 0.3, 0.45, 0.6, 0 (recovery), 0 (recovery 2)]%) and saved. Third, images were cropped to only include the fly’s body, tailored for each fly. Fourth, the Mean Square Error (MSE) between consecutive images across all pixels was calculated, giving one MSE value for each pair of consecutive frames at each isoflurane concentration *k%;*

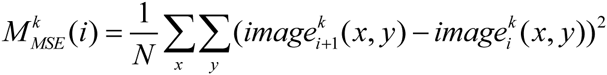

where N is the total number of pixels in each image, 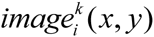 and 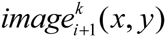 represent the grayscale value of pixel (x,y) in frame *i* and *i+1*, respectively, and the sum is taken over all pixels in the image. Finally, the resulting values were averaged over the two episodes of 6 air puffs in each block with isoflurane concentration *k* (^~^90 frames in total, Figure 4a) to obtain 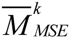. For comparison across flies, we further normalized the values for each fly by dividing the value in *k*% isoflurane 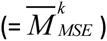 by the value in 0% isoflurane 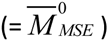. We refer to the resulting quantity as the Movement Index (MI^k^). MI values above and below 1.0 indicate increased and decreased movement compared to 0% isoflurane respectively. When computing MI for the recovery period we used the images from the last experimental block of each fly which was the 0% (recovery) for 8 flies and 0% (recovery 2) for 5 flies.

### Local Field Potential analysis

Electrophysiological data was recorded at 25 kHz and down-sampled to 1000 Hz for all subsequent analyses (Paulk et al., 2013). The most peripheral electrode site was removed from the analysis as it was outside the brain (see *Electrode Probe Insertion*). The remaining 15 electrodes sites were bipolar re-referenced by subtracting neighboring electrodes to obtain a set of 14 differential signals, which we refer to as channels hereafter (Figure 1c).

For SSVEP analysis we segmented the data in 2.3s epochs according to flicker configuration and isoflurane concentration. We removed line noise at 50Hz using the *rmlinesmovingwinc.m* function from the Chronux toolbox (http://chronux.org/, (Mitra and Bokil, 2007)) with 3 tapers, a window size of 0.7s and a step size of 0.35s.

#### Analyzing power

For each fly, we denote the power of the LFP during visual stimulation at frequency *ƒ*, in channel *i* (1-14), flicker configuration *l* (1-8) and isoflurane concentration *k%* ([0 (air), 0.06, 0.18, 0.3, 0.45, 0.6, 0 (recovery)]) as 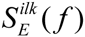 with subscript E meaning *evoked*. 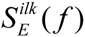 is in units of 10log10(μV^2^), averaged (in the log scale) over the 10 repetitions of the flicker configuration (see *Visual Stimuli*, Figure 4a). 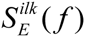 was calculated over the 2.3s trial period using the multi-taper method (*mtspectrumc.m*, http://chronux.org/, (Mitra and Bokil, 2007)) with three tapers, giving a half bandwidth of ^~^0.87Hz (Mitra and Pesaran, 1999), which is sufficiently fine for our claims in this paper. We denote spontaneous power at frequency *ƒ* in channel *i* and isoflurane *k%* as 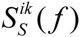 with subscript S meaning *spontaneous*. 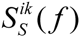 is the power averaged across four 2.3s long segments before the start of visual flicker presentation in units of 10log10(μV^2^) (Figure 4a).

When presenting results for k=0% (air), we corrected for baseline levels by subtracting the spontaneous power from SSVEP power

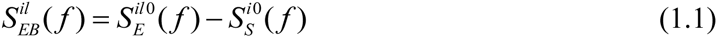

(subscript B for *baseline* correction). 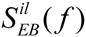 is reported in units of dB, emphasizing that the subtraction is performed after conversion to the log scale.

We use the symbols f_1_ and f_2_ to refer to the tag or twice the tag frequency respectively. The frequencies corresponding to f_1_ and f_2_ are flicker configuration dependent (e.g., when the flicker configuration was [13 13], f_1_=13Hz and f_2_=26Hz). We refer to power at frequency *ƒ*_n_ as the average power from −0.5 to +0.5 around the frequency of interest

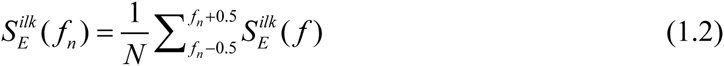

where N=4 is the number of frequency bins over which the sum is evaluated. The baseline corrected SSVEP power 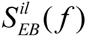 wasobtained by substitution of 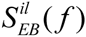 for 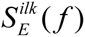 in equation (1.2).

When reporting SSVEP power for ipsilateral and contralateral flicker configurations, we separately averaged over the flicker configurations for each grouping and the corresponding tags (see *Grouping flicker configurations as ipsilateral or contralateral*, Figure 3a). For example, 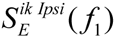 refers to the average power at f_1_ across the 6 flicker configurations where the flicker was presented ipsilateral to probe insertion site (13 Hz for [13 off], [13 13] and [13 17] and 17Hz for [17 off], [17 17] and [17 13]). The baseline corrected SSVEP power 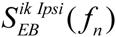 wasobtained by substituting 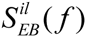 for 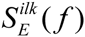 and repeating the derivation.

The effect of *k%* isoflurane on SSVEP power is denoted by the symbol Δ and obtained by subtracting respective values in 0% (air) isoflurane

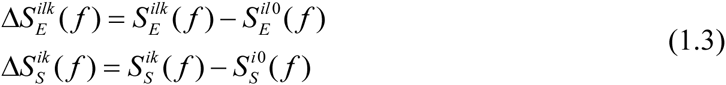

When reporting the effect on power at the tagged frequency (f=f_1_ or f_2_) we averaged the power around the tagged frequency as in equation (1.2). To obtain the average for ipsilateral/contralateral flicker configurations 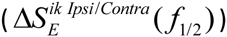, we repeated the derivation above with 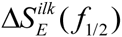 substituted for 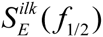.

#### Analyzing coherence

We analyzed coherence between channel pairs using the function *coherencyc.m* in the Chronux toolbox (Mitra and Bokil, 2007) with five tapers, giving a half bandwidth of 1.40Hz (Mitra and Pesaran, 1999), which is sufficient for our claims. Our notation and terminology for coherence parallel those used for reporting power, as described below.

SSVEP coherence for channel pair (*i,j*), flicker configuration *l* and isoflurane concentration *k%*, 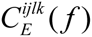 is calculated for the SSVEPs over the 2.3s trials and averaged over the 10 repetitions of the flicker configuration. As spontaneous coherence 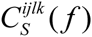 we report coherence averaged across four 2.3s long segments before the start of visual flicker presentation (Figure 4a).

Baseline corrected SSVEP coherence is used when presenting results in 0% (air) isoflurane and defined as

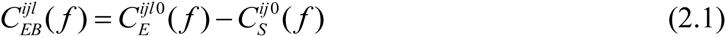

As for power, we refer to coherence at frequency 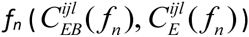 as the average coherence from −0.5 to +0.5 around the tagged frequency.

We calculated SSVEP and spontaneous coherence between all channel pairs, resulting in 91 (14*13/2) unique values at every frequency. To summarize these data in a concise way, we grouped channel pairs into periphery (channels 1-6) and center (channels 9-14) (Figure1b). We report periphery (P), center-periphery (CP), and center (C) coherence as averaged across all pairs of the electrodes within periphery, between center and periphery, and within center respectively;

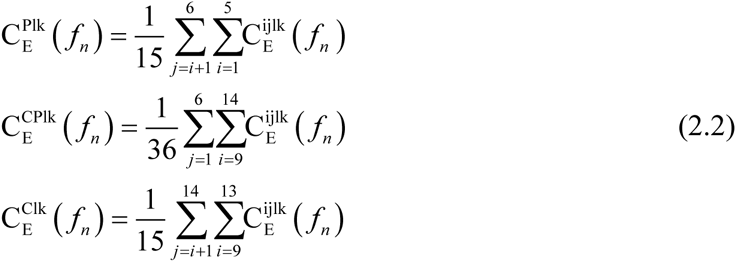

where the superscripts P, C and CP replace the channel pair superscript. The upper and lower limits of the sums reflect the grouping into peripheral (1-6) and center (9-14) channels and take into account that coherence is invariant with respect to channel order(C^ij^=C^ji^), while excluding coherence between a channel and itself (C^ii^=1). Our results were not sensitive to the exact grouping, such that other schemes, for example periphery = channels 2-5, center = channels 10-13, gave similar results. We obtained the analogous quantities for baseline corrected SSVEP coherence *ƒ*_*n*_, 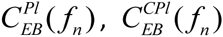 and 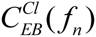 by substitution of 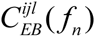 for 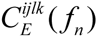 in equations (2.2).

Paralleling the power analysis, we report SSVEP coherence for ipsilateral and contralateral configurations at *ƒ*_1_, and *ƒ*_2_ (e.g. 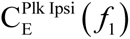 refers to the average coherence at f_1_ across 6 flicker conditions where the flicker was presented ipsilateral to probe insertion site, Figure 3a). The baseline corrected SSVEP coherence (e.g. 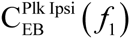) and the effect of k% isoflurane (e.g. 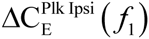) were defined similarly to the analogous quantities for power (e.g. 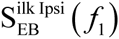 and 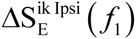

### Modeling the SSVEPs

We modelled the SSVEPs as the sum of two separate linear responses corresponding to the On and Off pathways (Figure 5a). The input (depicted as a square wave) is differentiated to extract points of luminance increments and decrements before splitting into two streams corresponding to the On and Off pathways. The responses of the On and Off pathways are summed to give the modelled SSVEP. Mathematically, the model’s output is given by

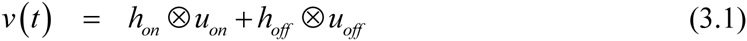

where *v* (*t*) is the output voltage, *h*_*on*_ and *h*_*off*_ are the impulse responses of the On and Off pathways respectively and ⊗ denotes convolution in the time domain. *u*_*on*_ and *u*_*off*_ are the half-wave rectified inputs to the On and Off pathways such that *u*_*on*_(*t*)=1 and *u*_*off*_(*t*)=1 signify an increase and decrease in luminance at time *t* respectively. Note that beyond the rectification nonlinearity the model is a linear multiple input/single output model (Bendat and Piersol, 2000). Our model assumes that the effect of isoflurane on the SSVEPs can be explained by changes to the impulse responses *h*_*on*_ and *h*_*off*_ alone.

We note that the nonlinearity is cancelled if the impulse response of the On pathway is identical and opposite to the impulse response of the Off pathway (Regan and Regan, 1988). Subbing *h*_*on*_ = –*h*_*off*_ into equation (3.1) and using *u*’(*t*) = *u*_*on*_(*t*)–*u*_*off*_(t), we obtain

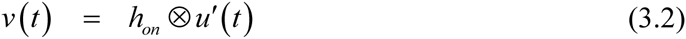

Because *u*’(*t*) is simply the (linearly) differentiated input (*u*’(*t*) = *u*(*t*)–*u*(*t*–1)), equation (3.2) shows that the model reduces to a single linear operation of the (linearly differentiated) input when *h*_*on*_ = –*h*_*off*_.

The model’s frequency response is given by

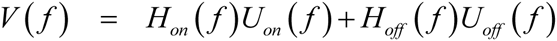

where the convolution in equation (3.1) is replaced by multiplication and capital letters represent the Fourier transforms of their respective variables. The power spectrum of the model’s response is given by

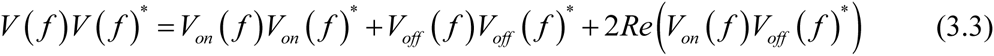

where we used *V*_*on*_(*ƒ*) = *H*_*on*_(*ƒ*)*U* _*on*_(*ƒ*) and *V*_*off*_(*ƒ*) = *H*_*off*_(*ƒ*)*U*_*off*_(*ƒ*) for the responses of the On and Off pathways to their respective inputs. *\** represents conjugation and *Re()* denotes taking the real part.

We note two things about equation (3.3). Firstly, the response at frequency *ƒ* is only a function of the responses of the On and Off pathways at frequency *ƒ*; there is no contribution from other frequencies. In the context of our experiments, this means that the model’s prediction of SSVEP power at *ƒ*_1_ and *ƒ*_2_ depends only on the stimulus, and the properties of the transfer functions (*H*_*on/off*_ (*ƒ*)) at *ƒ*_1_ and *ƒ*_2_.

Secondly, the model’s prediction for SSVEP power depends on the SSVEP power of the On (*V*_*on*_(*ƒ*)*V*_*on*_(*ƒ*)^*^) and Off pathways (*V*_*off*_(*ƒ*)*V*_*off*_(*ƒ*)^*^), but also on the cross spectrum *between* the responses (2*Re*(*V*_*on*_(*ƒ*)*V*_*off*_(*ƒ*)^*^)).

The On and Off impulse responses at each channel *i* and isoflurane concentration *k* 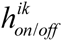 were estimated by averaging the LFP over 20 on-off cycles of the [1 1] flicker configuration, which we presented to three flies (see *Visual Stimuli*)(Figure 5b). Because the input is a square wave, the half-wave rectification effectively transforms the input into two pulse-trains (Figure 5a). We then used the estimated impulse responses at each channel and isoflurane concentration together with equation (3.1) to predict SSVEPs for the [13 13] and [17 17] flicker configurations by setting the input to a 50% duty-cycle square wave with periods (1/13Hz) and (1/17Hz) respectively. By computing the Fourier transform of the modelled SSVEPs at each channel *i*, the two flicker configurations *l* ([13 13] and [17 17]) and isoflurane concentration *k*, we obtained the model’s prediction for SSVEP power 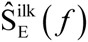. The model’s prediction for the effect of *k*% isoflurane 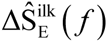 is obtained by substituting 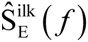 for the measured SSVEP power 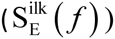 in equation (1.3). The quantities 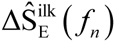 and 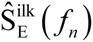 were obtained by averaging from −0.5 to +0.5 around the frequency of interest. Note that the model does not have any degrees of freedom for fitting, once the impulse responses are determined by the simple averaging of the data over 20 on-off cycles of the [1 1] flicker configuration. Nothing further is estimated or fit from the data.

#### Evaluating the SSVEP model

To investigate the relationship between the model and data we performed linear regression between the model-predicted and observed effect of isoflurane on SSVEP power

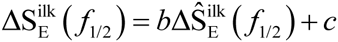

b and c were estimated in R (https://www.r-project.org/, R Core Development Team, 2015) using the *lm* function. We report the Pearson’s correlation coefficient ρ between the model’s prediction and the observed data and 95% confidence intervals on the slope (b) and intersect (c) obtained by the *confint* function. Note that a perfect fit between model and data is given by ρ = 1 and the line (b=1, c=0).

We performed the regression over three flies (3), all channels (1-14), two flicker configurations ([13 13] and [17 17]) and both f_1_ and f_2_, giving 168 paired data points in total. We used the highest concentration of isoflurane (represented as H in superscript of [delta S^H^_E_ in Figure 5g]) presented to each fly, k=0.6% for two flies and k=0.45% for one fly (see *Isoflurane delivery*).

### SNR-based estimation of coherence

To investigate the relationship between evoked power and coherence we assumed a linear framework in which the SSVEPs for each channel pair are related through a linear transfer function in the presence of noise (Figure 6b). This framework is conceptually related to the SSVEP model but completely independent in its implementation and evaluation.

In this framework, the SSVEP at channel *i*, given by v_i_(t) passes through the linear transfer function H_ij_(f) to give the SSVEP at channel *j*, v_j_(t). Independent noise enters at each channel separately (n_i_(t) and n_j_(t)) to give the recorded responses y_i_(t) and y_j_(t). Under these assumptions, squared coherence between channel pairs has an analytical description (for the detailed derivation, see (Bendat and Piersol, 2000) for example);

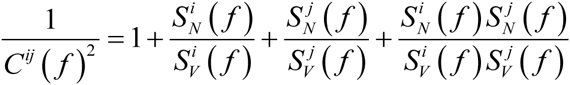

where 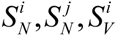 and 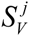 are the power spectrums of n_i_, n_j_, vi and v_j_, respectively. If we define the Signal to Noise Ratios (SNRs) 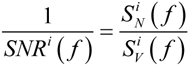 and 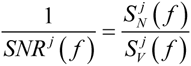 then

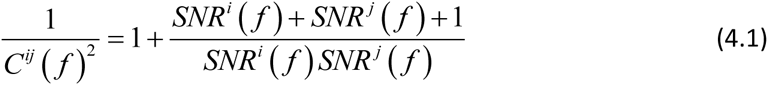

Thus, in this simplified setting coherence is totally determined by the SNRs at the respective channels.

To evaluate the SNR-based coherence estimate we quantified the SNR at each channel and used equation (4.1) to obtain the model’s prediction of SSVEP coherence. First, we recalculated 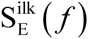 using the same number of tapers used for the coherence analysis (i.e., 5 tapers, see *Local Field Potential analysis*). We then fitted power law noise to the observed SSVEP power 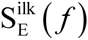

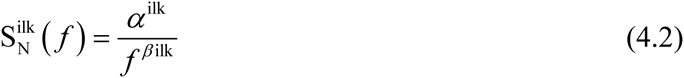

where we excluded f values from −1.4 to +1.4Hz around f_1_ and f_2_ for each flicker configuration (1.4Hz corresponds to the half-bandwidth for the coherence measurement, see *Local Field Potential analysis*). The purpose of the fit is to estimate the level of neural activity that is not directly tagged by visual flickers. Following the convention in the SSVEP literature (Norcia et al., 2015), we considered the non-tagged activity representing the level of noise (e.g., n(t) or S_N_(f)). The parameters 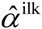 and 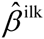 were estimated by linear regression in the log-log scale in the range 1-50Hz and used to define the noise spectrum.

We define the SNR at frequency f as the observed SSVEP power at f divided by the interpolated noise spectrum at f

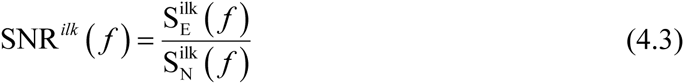

An example of the estimated noise spectrums and resulting SNRs for two exemplar channels is shown in Figure 6c-d. Figure 6e shows the resulting coherence estimate.

The predicted effect of *k%* isoflurane on coherence for channel pair (*i,j*) is obtained by subtracting the predicted coherence in 0% (air) isoflurane from the predicted coherence in *k%* isoflurane

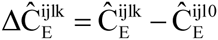

For the further analyses, we grouped the electrode pairs into the periphery, center and center-periphery and for ipsilateral and contralateral flicker configurations separately as described before (presented in Figure 6f).

To provide an overall measure of fit for the SNR-based estimation of coherence we calculated the Mean Square Error (MSE) between the model’s prediction and the observed effect of 0.6% isoflurane on SSVEP coherence in each fly, across all channel pairs (91), flicker configurations (8) and f_1_ and f_2_

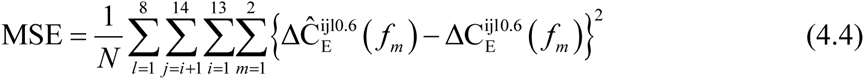

where N is the total number of terms in the sum, N=91*8*2=1456 (for each of thirteen flies).

#### Separating the contribution of “noise” and “signal” to the SNR-based estimation of coherence

The SNR-based estimation of coherence is completely determined by the SNR (equation (4.1)), which in turn is a function of the estimated noise levels from non-tagged frequency as well as the observed SSVEP power at the tagged frequency (equation (4.3)). To isolate the contribution of the noise and SSVEP power to the coherence estimate, we defined two additional variants of SNR. In the first, SNR_FN_, (subscript FN for *fixed noise*) we fixed the noise spectrum to that fitted in 0% (air) isoflurane

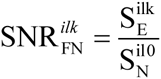

thereby removing the influence of isoflurane on noise levels. In the second, SNR_FE_ (subscript FE for *fixed SSVEP* power) we fixed the SSVEP power to that observed in 0% (air) isoflurane

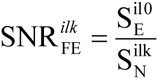

thereby removing the influence of isoflurane on SSVEP power. 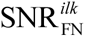 and 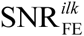 are used together with equation (4.1) to obtain two additional estimates of coherence (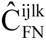 and 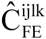), the effects of *k%* isoflurane (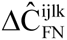 and 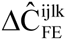), as well as the grouped coherence over electrode pairs and flicker configurations as described before (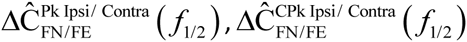 and 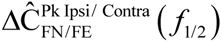, presented in Figure 6g-h). Paired t-tests between the MSEs (equation (4.4), obtained for each fly separately) between the observed effects of isoflurane on coherence and those predicted by each SNR variant were used for assessing statistical significance.

### Statistical analysis

We used R (https://www.r-project.org/, R Core Development Team, 2015) and lme4 (Bates et al., 2015) to perform linear mixed effect analysis of the data. Throughout, the response variable is either power 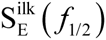 or coherence 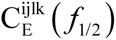 with four factors, *isoflurane, channel location, harmonic*, and *fly*.

As to the factors *flicker configuration* and *response frequency*, only a subset of all combinations are relevant for our claims. Specifically, when the flicker configuration was [13 13], [13 off] or [off 13] we analyzed power at f_1_=13Hz and f_2_=26Hz. When the flicker configuration was [17 17], [17 off] or [off 17] we analyzed power at f_1_=17Hz and f_2_=34Hz. When the flicker configuration was [13 17] or [17 13] we analyzed power at f_1_=13Hz, 17Hz, f_2_=26Hz and 34Hz. Thus, replacing the *response frequency* factor, we included the factor *flicker location* (categorical, 2 level), that corresponds to the division into ipsi-and contralateral flicker configurations (see Figure 3a and *Grouping flicker configurations as ipsilateral and contralateral*).

Among those factors, we focused on the crucial isoflurane specific effects by including interactions between *isoflurane* and *flicker location, isoflurane* and *harmonic* and *isoflurane* and *channel location*, as well as the triple interaction between *isoflurane, channel location* and *harmonic*. In addition, our results in 0% (air) isoflurane imply harmonic dependent effects for *flicker location* and *channel location* so we included interactions between *flicker location* and *harmonic*, and *channel location* and *harmonic*. We included random intercepts for all fixed factors to account for possible correlation between the levels of each factor.

To test for the effect of a given factor or interaction we performed likelihood ratio tests between the full model described above and a reduced model without the factor or interaction in question (Bates et al., 2015). When applicable, we adjusted p-values using the false discovery rate (FDR, (Yekutieli and Benjamini, 1999))

## Acknowledgments

We would like to thank Angelique C Paulk for helpful discussion and assistance using the multi-electrode preparation.

NT was funded by ARC Future Fellowship (FT120100619) and Discovery Project (DP130100194).

BVS was funded by NHMRC Project APP1103923

## References

Alkire, M. T., Hudetz, A. G. & Tononi, G. 2008. Consciousness and anesthesia. Science (New York, N.Y.), 322, 876–880.10.1126/science.1149213

Bates, D., Mächler, M., Bolker, B. & Walker, S. 2015. Fitting Linear Mixed-Effects Models using lme4. Journal of Statistical Software, 67.10.18637/jss.v067.i01

Bendat, J. S. & Piersol, A. G. 2000. Random Data: Analysis and Measurement Procedures, New York, John Wiley & Sons, Inc.

Borst, A. & Euler, T. 2011. Seeing Things in Motion: Models, Circuits, and Mechanisms. Neuron, 71.10.1016/j.neuron.2011.08.031

Brown, E. N., Purdon, P. L. & Van Dort, C. J. 2011. General anesthesia and altered states of arousal: a systems neuroscience analysis. Annu Rev Neurosci, 34, 601–28.10.1146/annurev-neuro-060909-153200

Capilla, A., Pazo-Alvarez, P., Darriba, A., Campo, P. & Gross, J. 2011. Steady-State Visual Evoked Potentials Can Be Explained by Temporal Superposition of Transient Event-Related Responses. PLoS ONE, 6.10.1371/journal.pone.0014543

Casali, A. G., Gosseries, O., Rosanova, M., Boly, M., Sarasso, S., Casali, K. R., Casarotto, S., Bruno, A. M., Laureys, S., Tononi, G. & Massimini, M. 2013. A Theoretically Based Index of Consciousness Independent of Sensory Processing and Behavior. Science Translational Medicine, 5.10.1126/scitranslmed.3006294

Cimenser, A., Purdon, P. L., Pierce, E. T., Walsh, J. L., Salazar-Gomez, A. F., Harrell, P. G., Tavares-Stoeckel, C., Habeeb, K. & Brown, E. N. 2011. Tracking brain states under general anesthesia by using global coherence analysis. Proceedings of the National Academy of Sciences, 108, 8832–8837.10.1073/pnas.1017041108

Egelhaaf, M. & Borst, A. 1989. Transient and steady-state response properties of movement detectors. Journal of the Optical Society of America. A, Optics and image science, 6, 116–127.10.1364/JOSAA.6.000116

Ferrarelli, F., Massimini, M., Sarasso, S., Casali, A., Riedner, B. A., Angelini, G., Tononi, G. & Pearce, R. A. 2010. Breakdown in cortical effective connectivity during midazolam-induced loss of consciousness. Proceedings of the National Academy of Sciences, 107, 2681–2686.10.1073/pnas.0913008107

Franks, N. P. 2008. General anaesthesia: from molecular targets to neuronal pathways of sleep and arousal. Nature Reviews Neuroscience, 9, 370–386.10.1038/nrn2372

Garcia, P. S., Kolesky, S. E. & Jenkins, A. 2010. General Anesthetic Actions on GABAA Receptors. Current Neuropharmacology, 8, 2–9.10.2174/157015910790909502

Haag, J. & Borst, A. 2008. Electrical Coupling of Lobula Plate Tangential Cells to a Heterolateral Motion-Sensitive Neuron in the Fly. The Journal of Neuroscience, 28, 14435–14442.10.1523/JNEUROSCI.3603-08.2008

Joesch, M., Schnell, B., Raghu, S., Reiff, D. F. & Borst, A. 2010. ON and OFF pathways in Drosophila motion vision. Nature, 468, 300–304.10.1038/nature09545

Kirszenblat, L. & Van Swinderen, B. 2015. The Yin and Yang of Sleep and Attention. Trends in Neurosciences, 38, 776–786.10.1016/j.tins.2015.10.001

Kottler, B., Bao, H., Zalucki, O., Imlach, W., Troup, M., Van Alphen, B., Paulk, A., Zhang, B. & Van Swinderen, B. 2013. A Sleep/Wake Circuit Controls Isoflurane Sensitivity in Drosophila. Current Biology, 23.10.1016/j.cub.2013.02.021

Lee, U., Blain-Moraes, S. & Mashour, G. A. 2015. Assessing levels of consciousness with symbolic analysis. Philosophical Transactions of the Royal Society of London A: Mathematical, Physical and Engineering Sciences, 373, 20140117.10.1098/rsta.2014.0117

Lee, U., Ku, S., Noh, G., Baek, S., Choi, B. & Mashour, G. A. 2013. Disruption of frontal-parietal communication by ketamine, propofol, and sevoflurane. Anesthesiology, 118, 1264–75.10.1097/ALN.0b013e31829103f5

Lewis, L. D., Weiner, V. S., Mukamel, E. A., Donoghue, J. A., Eskandar, E. N., Madsen, J. R., Anderson, W. S., Hochberg, L. R., Cash, S. S., Brown, E. N. & Purdon, P. L. 2012. Rapid fragmentation of neuronal networks at the onset of propofol-induced unconsciousness. Proceedings of the National Academy of Sciences, 109.10.1073/pnas.1210907109

Liu, X., Lauer, K. K., Ward, B. D., Rao, S. M., Li, S. J. & Hudetz, A. G. 2012. Propofol disrupts functional interactions between sensory and high - order processing of auditory verbal memory. Human Brain Mapping, 33, 2487–2498.10.1002/hbm.21385

Mashour, G. A. 2013. Cognitive unbinding: a neuroscientific paradigm of general anesthesia and related states of unconsciousness. Neuroscience and biobehavioral reviews, 37, 2751–2759.10.1016/j.neubiorev.2013.09.009

Mashour, G. A. 2014. Top-down mechanisms of anesthetic-induced unconsciousness. Frontiers in systems neuroscience, 8, 115.10.3389/fnsys.2014.00115

Mitra, P. & Bokil, H. 2007. Observed brain dynamics, Oxford University Press

Mitra, P. P. & Pesaran, B. 1999. Analysis of dynamic brain imaging data. Biophysical journal, 76, 691–708.10.1016/S0006-3495(99)77236-X

Murphy, M., Bruno, M.-A., Riedner, B. A., Boveroux, P., Noirhomme, Q., Landsness, E. C., Brichant, J.-F., Phillips, C., Massimini, M., Laureys, S., Tononi, G. & Boly, M. 2011. Propofol anesthesia and sleep: a high-density EEG study. Sleep, 34, 283

Norcia, A. M., Appelbaum, L. G. & Ales, J. M. 2015. The steady-state visual evoked potential in vision research: A review. Journal of vision, 15.10.1167/15.6.4

Paulk, A. C., Kirszenblat, L., Zhou, Y. & Van SWINDEREN, B. 2015. Closed-Loop Behavioral Control Increases Coherence in the Fly Brain. The Journal of Neuroscience, 35, 10304–10315.10.1523/JNEUROSCI.0691-15.2015

Paulk, A. C., Zhou, Y., Stratton, P., Liu, L. & Van SWINDEREN, B. 2013. Multichannel brain recordings in behaving Drosophila reveal oscillatory activity and local coherence in response to sensory stimulation and circuit activation. Journal of neurophysiology, 110, 1703–1721.10.1152/jn.00414.2013

Regan, M. P. & Regan, D. 1988. A frequency domain technique for characterizing nonlinearities in biological systems. Journal of Theoretical Biology, 133, 293–317.10.1016/S0022-5193(88)80323-0

Reiff, D. F., Plett, J., Mank, M., Griesbeck, O. & Borst, A. 2010. Visualizing retinotopic half-wave rectified input to the motion detection circuitry of Drosophila. Nature Neuroscience, 13, 973–978.10.1038/nn.2595

Reisenman, C., Haag, J. & Borst, A. 2003. Adaptation of response transients in fly motion vision. I: Experiments. Vision Research, 43, 1293–1309.10.1016/S0042-6989(03)00091-9

Sarasso, S., Boly, M., Napolitani, M., Gosseries, O., Charland-VERVILLE, V., Casarotto, S., Rosanova, M., Casali, A., Brichant, J.-F., Boveroux, P., Rex, S., Tononi, G., Laureys, S. & Massimini, M. 2015. Consciousness and Complexity during Unresponsiveness Induced by Propofol, Xenon, and Ketamine. Current Biology, 25, 3099–3105.10.1016/j.cub.2015.10.014

Sarasso, S., Rosanova, M., Casali, A. G., Casarotto, S., Fecchio, M., Boly, M., Gosseries, O., Tononi, G., Laureys, S. & Massimini, M. 2014. Quantifying cortical EEG responses to TMS in (un)consciousness. Clinical EEG and neuroscience, 45, 40–49.10.1177/1550059413513723

Supp, G. G., Siegel, M., Hipp, J. F. & Engel, A. K. 2011. Cortical hypersynchrony predicts breakdown of sensory processing during loss of consciousness. Current biology: CB, 21, 1988–1993.10.1016/j.cub.2011.10.017

Van Swinderen, B. 2006. A succession of anesthetic endpoints in the Drosophila brain. J Neurobiol, 66, 1195–211.10.1002/neu.20300

Van Swinderen, B. & Kottler, B. 2014. Explaining general anesthesia: a two step hypothesis linking sleep circuits and the synaptic release machinery. BioEssays, 36, 372–381.10.1002/bies.201300154

Yekutieli, D. & Benjamini, Y. 1999. Resampling-based false discovery rate controlling multiple test procedures for correlated test statistics. Journal of Statistical Planning and Inference, 82, 171–196.10.1016/S0378-3758(99)00041-5

Zalucki, O. H., Menon, H., Kottler, B., Faville, R. & Day, R. 2015. Syntaxin1A-mediated Resistance and Hypersensitivity to Isoflurane in Drosophila melanogaster. Anesthesiology.

